# Algorithmic construction of topologically complex biomineral lattices via cellular syncytia

**DOI:** 10.1101/2024.02.20.580924

**Authors:** Pranav Vyas, Charlotte Brannon, Laurent Formery, Christopher J. Lowe, Manu Prakash

## Abstract

Biomineralization is ubiquitous in both unicellular and multicellular living systems [1, 2] and has remained elusive due to a limited understanding of physicochemical and biomolecular processes [3]. Echinoderms, identified with diverse architectures of calcite-based structures in the dermis[4], present an enigma of how cellular processes control shape and form of individual structures. Specifically, in holothurians (sea cucumbers), multi-cellular clusters construct discrete single-crystal calcite ‘ossicles’ (*∼* 100 *µ*m length scale), with diverse morphologies both across species and even within an individual animal [5]. The local rules that might encode these unique morphologies in calcite ossicles in holothurians remain largely unknown. Here we show how transport processes in a cellular syncytium impart a top-down control on ossicle geometry via symmetry breaking, branching, and fusion in finite cellular clusters. As a unique example of cellular masonary, we show how coordination within a small cluster of cells builds calcite structures about an order of magnitude larger than any individual participating cell. We establish live imaging of ossicle growth in *Apostichopus parvimensis* juveniles revealing how individual crystalline seeds (*∼* 1 *−* 2 *µ*m) grow inside a multi-cellular syncytial complex with the biomineral completely wrapped within a membrane-bound cytoplasmic sheath. Constructing a topological description of ossicle geometries from 3D micro-CT (computational tomography) data reveals the hidden growth history and conserved patterns across ossicle types. We further demonstrate vesicle transport on the surface of the ossicle, rather than cell motility, regulates material transport to the ossicle tips via a unique cytoskeletal architecture. Finally, using reduced order models of conserved transport on self-closing active branching networks, we highlight the hidden universality in the growth process of distinct ossicles. The system presented here serves as a unique playground merging top-down cellular physiology and classical branching morphogenesis [6] with bottom-up non-equilibrium mineralization [7] processes at the interface of living and non-living matter [8].

## Introduction

Biomineralization has evolved multiple times independently [1, 9] across the tree of life encompassing both unicellular and multicellular organisms. The study of biomineralized structures across numerous systems has enabled discovery of diverse functions including magnetotaxis [10, 11], gravity sensing [12, 13], vision [14, 15], predator defense [16, 17], mineral regulation and detoxification [18, 19] and structural support [1, 20, 21, 2]. However, how morphological complexity of mineralized structures is encoded has remained an open question for over a century [22, 23].

Biomineralization in metazoans relies on cellular processes that sequester inorganic mineral ions and specialized proteins to privileged spaces, facilitating stable mineral growth [2, 24]. To build functional and precise geometry out of a crystalline material, processes across multiple spatio-temporal scales must be integrated: crystallization at the molecular scale, regulation and transport of precursor materials at the cellular scale, resource allocation and signaling at the tissue scale, and developmental patterning at the organism scale. It is remarkable that not only single cells [25], but multi-cellular clusters are also capable of coordinating activity across scales to build mineralized structures significantly larger than individual cells [2].

In the past, several systems including cnidarians, mollusks, and echinoderms have been utilized to reveal the bio-molecular and mineral phase behavior during biomineralization [9, 26]. Amongst these, echinoderms stand out due to the large scale occurrence of biomineralization across their dermal tissue in the form of numerous fused or discrete units called ossicles which provide skeletal support [27]. In particular, the skeletal spicule of larval sea urchin has proven to be instrumental in developing insights into biomineralization gene regulatory networks (GRNs) [28, 29], developmental programs [30, 31], cellular processes [32, 33], material transport processes [34, 35], and aspects of mineral phase change [36]. In comparison, very little is known about fundamental processes that generate the vast diversity of ossicle shapes and forms seen in holothurians [37].

Holothurians, commonly known as sea cucumbers, distinguish themselves amongst echinoderms with their soft and flexible bodies. This occurs both due to much smaller evolved sizes of ossicles (*∼* 100*µm* in size) and lack of fusion between them to form a contiguous skeleton, which is characteristically observed in other adult echinoderms [4]. This distinction has been postulated to occur due to an identified evolutionary regression of genes downstream of biomineralization signaling pathways conserved across echinoderms [38]. A single adult holothurian can contain more than 20 million discrete ossicles [39] that are distributed across the entire body wall. Across roughly 1800 distinct sea cucumber species, more than 50 morphotypes of ossicle shapes have been described with descriptive shapes including tables, wheels, holed plates, anchors, spikes which are widely utilized in classification of both extant and extinct taxa [5], making these structures important from a palaeontological perspective as well. Ossicles are highly complex structures, both geometrically as well as topologically, due to their multi-genus(multi-holed) shapes. This approach of micro-scale skeletonization across the animal body is thought to have evolved independently in sponges, holothurians, corals, flatworms, tunicates, and several mollusks, which has allowed distinct cell-tissue scale approaches to be utilized for similar discretized growth of complex shapes [40]. However, how multi-cellular processes regulate growth of such a dramatic diversity of complex mineralized structur es remains largely unknown.

Several studies of the holothurian body wall have provided insights into the local biomineralization niche created by the participating cells and snapshots of multiple ossicles at different growth stages [41, 42]. However, continuous live imaging of single ossicle growth has never been possible. Ossicles vary in shape and form not only across species, but even within a single individual. This morphological diversity within a single individual allows us to observe simultaneous iterations of mineral growth, enabling us to study multicellular coordination of ossicle construction, the spatial scale and fidelity of shape programming, the universal biophysical mechanisms of growth, and the overlaying layers of biological control that give rise to the ossicle diversity.

The near transparent nature of early juvenile stages (Figure 1A), makes the interface between living and non-living phases of mineralized holothurian tissues clearly visible. Thus, in this study, we utilize the early juvenile stage of sea cucumber *Apostichopus parvimensis* (widely present in Monterey Bay, California) to directly observe ossicle morphogenesis. The metamorphosis of the pelagic larval form of the animal into its benthic (Figure S1) juvenile form (*∼* 20 *−* 40 days post fertilization(dpf)) is accompanied by the simultaneous growth of multiple ossicles throughout the animal body (Figure 1A,B). The chemical composition of holothurian ossicles has been previously determined to be magnesium-rich calcite (crystalline polymorph of CaCO_3_) and organic matrix molecules [43, 34] which is similar to other echinoderms [44].

**Figure 1:**
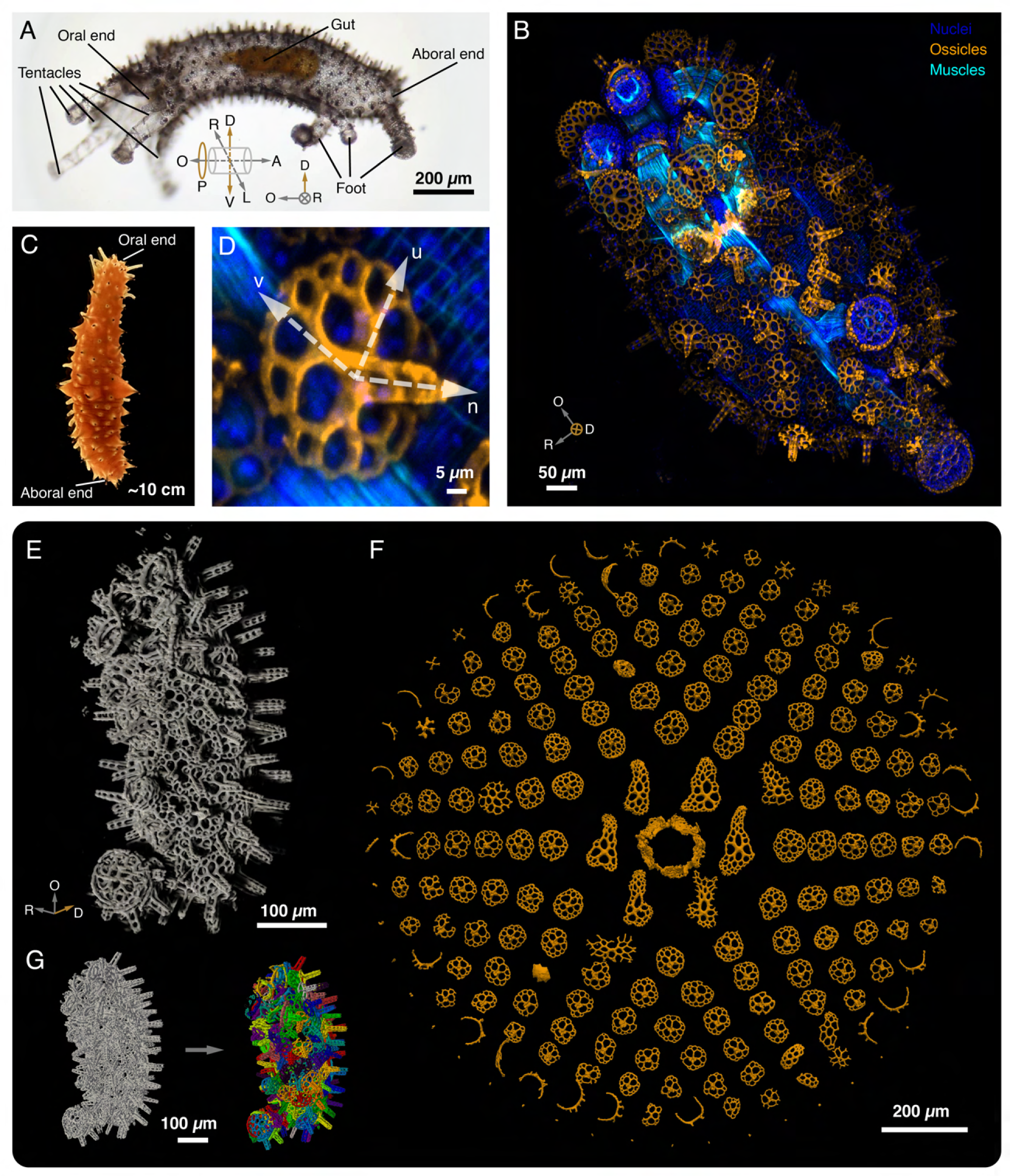
Finite and diverse biomineral structures in juvenile *Apostichopus parvimensis*. (A) At 20 40 dpf the pelagic (Auricularia) larval form of the animal metamorphoses into its benthic (Pentacula) juvenile form. This is accompanied by the simultaneous growth of multiple magnesium-rich calcite ossicles throughout the animal body. The cylindrical marker represents the body plan axes within the animal. The animal displays a pentaradial symmetry with an overlayed bilateral symmetry. O - oral end, A - aboral end, D - dorsal surface, V - ventral surface, L - left side, R- right side, P - pentaradial symmetry. (B) Maximum intensity projection for a whole mount confocal scan of a fixed juvenile *A. parvimensis*. Cell nuclei are labeled with Hoechst (dark blue), muscles are labeled with phalloidin (cyan), and ossicles are labeled with calcein (gold). The animal displays a diversity of ossicle geometries distributed according to body plan symmetries or organ-specific niches. (C) A 1-2 year-old adult *A. parvimensis* displaying an opaque body due to high pigmentation throughout the dermal tissue. (D) Enlarged view of a single ossicle from panel B. Each ossicle is an independent structure growing just below the outer epithelium that never merges with other nearby growing ossicles. Most of the structure grows planarly with respect to the local epithelial surface, though some ossicles possess vertical spines as well. U,V - locally coplanar axes to the body surface, N - normal to the body surface. (E) Micro-CT scans of fixed whole juveniles enable 3D visualization of ossicle structures at various stages of development. The lower row depicts a representative figure for the segmentation of the volumetric raw dataset into meshes representing individual ossicles. (F) All ossicles (total 213) from an individual, are segmented, aligned, and rearranged based on size (decreasing radially outward) to represent the morphological diversity and scales present in a single animal.

We study ossicle morphogensis in this system by quantitatively describing ossicle geometry and topology using high-resolution 3D micro-CT datasets and utilizing live imaging to decipher the growth algorithm followed by each ossicle. To understand how cell clusters control ossicle growth, we perform TEM and confocal imaging and show that ossicle growth happens within an ultra-thin syncytial niche throughout its entire development. We discover a feedback between cell shape and ossicle shape, where cellular control of mineral deposition controls ossicle shape while a tight syncytial wrapping forces the cellular cluster to assume the shape of the ossicle it is building. The accretive nature of growth and rigidity of the biomineral enforce immutability of geometry during the ossicle growth trajectory. Any variations or errors are thus literally ‘set in stone’ over the course of growth, leading to a history dependent evolution of the lattice structure. Any variations in the geometry established during the early phases of growth can thus dramatically alter the topology generated during the later stages. Through live imaging of this niche, we discover active processes overlaying ossicle geometry, controlling material transport and directional growth. Finally, using a simple two-dimensional constrained branching growth model we demonstrate observed emergent universal features of ossicle morphogenesis. Taken together, our findings reveal coordination of multiscale processes such as filopodial search and vesicle transport to construct topologically complex mineralized structures through branching morphogenesis.

## Results

### Holothurian ossicles are finite, disordered lattice displaying geometrical and topological variations

Juvenile holothurians display a distributed skeleton of calcite ossicles anchored between the dermal tissue matrix and the epithelium across the body (Figure 1A, 1B). To visualize the ossicles, animals were cultured in the presence of calcein, a fluorescent calcium chelator, resulting in the fluorescent labeling of all mineral structures [45] (see methods for details)(Figure 1B). Employing confocal microscopy, the ossicles can be identified as distinct perforated plates (as shown in an enlarged view in Figure 1D), exhibiting a striking variability in morphology throughout different parts of the body.”

To study ossicle morphologies in three-dimensions, we performed high-resolution micro-CT (min *∼* 0.8*µm* voxel size) scans on whole juveniles at different developmental stages to generate the first comprehensive database of spatially tagged 3D point clouds of individual ossicles (see Methods)(Figure 1E). Figure 1F depicts all ossicles from a single animal aligned planarly and sorted by size, showcasing the diversity of forms including variation in shape, size, and symmetry. Despite this diversity, the ossicles appear to be constructed out of very similar structural motifs.

Access to the exact geometry and spatial location of every ossicle (Figure 2A, 2B) allows us to examine the variation of ossicle shape in conjunction with the animal’s body plan. We observe 11 distinct ossicle types across the body: pillared tables, front plates, rear plates, foot pad plates, foot side plates, elongated plates, c-rods, larval ossicle, a basket cluster, the internal calcareous ring and several minuscule ossicles, some of which are previously described in taxonomic literature [5] (Figure 2B). One of the most common architectures is the pillared table, covering *∼* 75% of animal mineral volume and *∼* 70% of the total ossicle count (*∼* 213 ossicles). These morphological distinctions tend to be correlated with the distinct tissue niches present within the animal, hinting towards their functional roles. For example, the arrangement of front plates obeys the pentaradial symmetry along the oral-aboral axis (Figure 2A, B) supporting the oral opening; c-rods are localized within tentacles, possibly reinforcing the tubes against circumferential loads; and the flat foot plates are localized on the foot surface without any protruding pillar growth, reinforcing the structure of the foot. At the same time, distinct ossicle morphologies are also observed growing close to each other in similar niches, hinting towards a lack of a unique prescription of shape from local microenvironments (Figure S2B). Because of their comparable planar geometries, we will henceforth focus on base geometry of pillared tables, head/rear and side flat plates (together referred to as ‘other plates’), and the foot pad plate.

**Figure 2:**
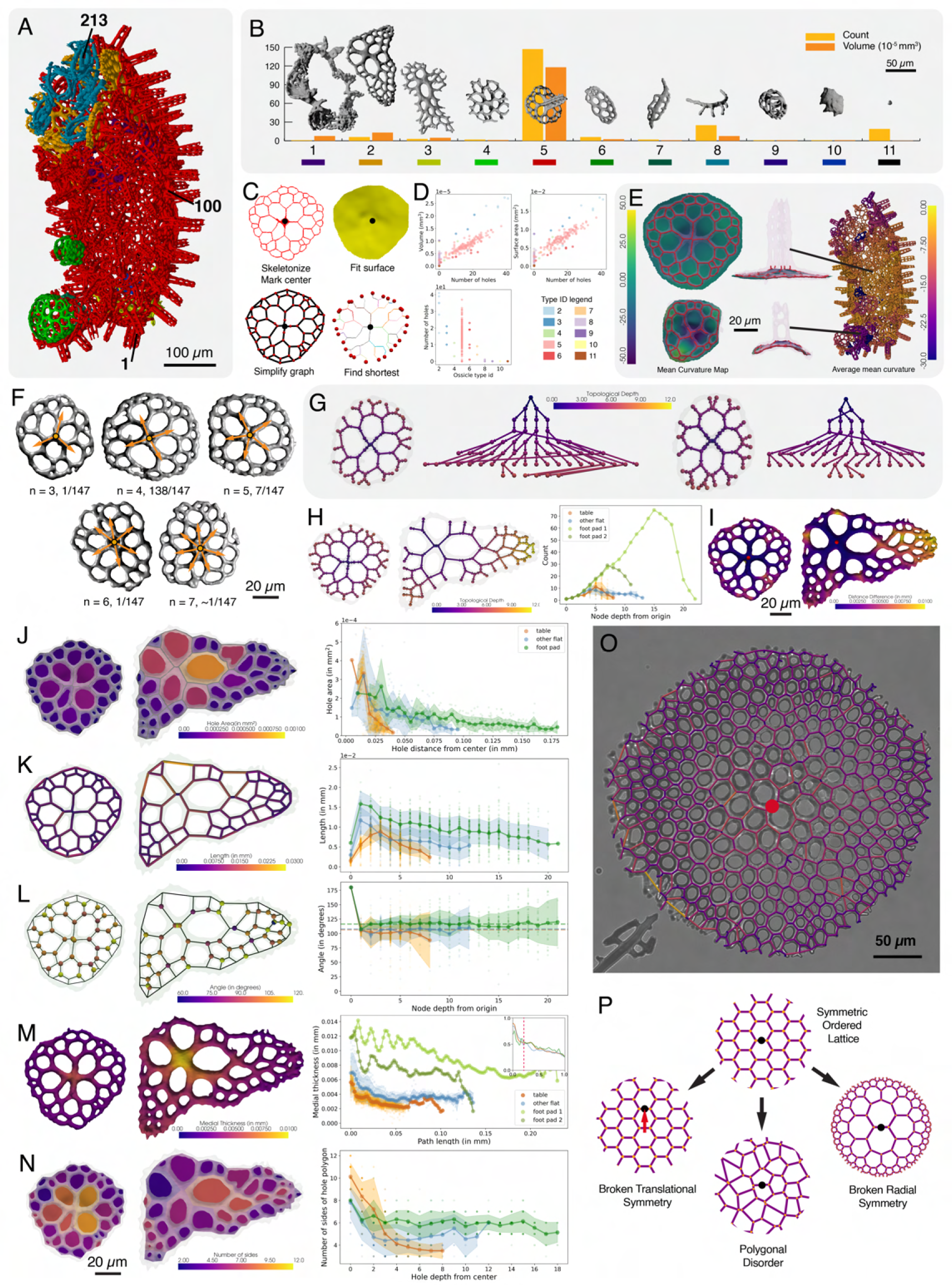
Developmental, geometrical and topological description of ossicles from juvenile *A. parvimensis*. (A) Segmented Micro-CT data with color labels highlighting positions of developmentally distinguishable ossicle types. (B) Ossicle type distribution with volume and number fraction data from a single juvenile. Legend: 1 - internal calcareous ring, 2 - front plates, 3 - rear plates, 4 - foot pad plates, 5 - pillared tables, 6 - foot side plates, 7 - elongated plates, 8 - c-rods, 9 - basket cluster, 10 - larval ossicle, 11 - minuscule ossicles. Table ossicles compose most of the mineral volume of the animal both in terms of volume (75%) as well as count (70%). (C) Segmented volumes of ossicles are converted into meshes, which can then be skeletonized (top left). The center of the ossicle is identified as the mid-point of the thickest edge, or most symmetric centrally located site. The skeletons are simplified into discrete graphs (bottom left) and surfaces representing the base of table ossicles (top right) can be generated through fitting one the graphs. Additional topological and geometrical characteristics are also extracted using the graphs, such as boundary nodes, and minimal paths connecting to the center (bottom right). (D) Scaling of ossicle surface area and volume with the number of holes demonstrating addition of more holes as the dominant mode of size increase (top left and top right). Hole size variation across ossicle types in a single individual (bottom left). (E) Left column demonstrates variation of mean curvature field within a surface fit to the base of the pillared table ossicles. Whole animal plot demonstrating variation in the base curvature of table ossicles depending on the location on the animal body (right panel). The ossicles conform to the local curvature of the body, showing higher curvature near the foot. (F) Pillared table ossicles displaying a range of central symmetries, varying from three to seven branches, with four pronged symmetry being the most abundant. (G) Comparison of topologies of two pillared tables with identical central symmetries highlights reduced conservation with increasing node depth away from the center. The arrows connect nodes at each edge, and point from a node with lower node depth towards one with higher node depth. (H) Left panel shows a comparison between similar topological graphs for pillared table and front plate ossicles. Right panel shows a comparison of number of nodes at a given node depth from the center, across pillared tables, other flat plates, and foot pad ossicles. The peak number of nodes increase almost linearly for foot pad ossicles, after which it decays due to irregularities at the boundaries. For variation also increases with node depth highlighting conservation of topology near the center. (I) A color map displaying the difference between geodesic vs radial distance for every point on the ossicle mesh. More distal and asymmetrically placed regions display higher difference between the two. (J-N) Color maps for distinct ossicle measurements along with combined statistics across distinct ossicles. (J) Meshes representing holes are extracted from base surface mesh. The hole areas decrease with distance from center. The decrease rate is more gradual for larger ossicles. (K) The branch lengths measured as distance between connected nodes, also display a decreasing trend with radial position. The variations first increase, right near the center, but then remain constant across the structure. (L) The branch bifurcation angles, measured only at nodes generating two nodes with higher depth, remain nearly conserved node depth value. The variation in observed angles however increases with node depth. The mean angle values are 106.5*^o^* for pillared tables, 107.7*^o^* for other plates, and 116.5*^o^* for foot pad ossicle - approaching that for the regular hexagonal lattice (120*^o^*) with increasing size. (M) The ossicle meshes displaying local medial thickness calculated on the nearest point on the skeleton of the mesh and averaged over a kernel to display smooth thickness gradients. The right panel shows medial thickness (t) with respect to the shortest path length distance (d) from the center node along the branches of the skeleton. The inset shows plot of mean values for different ossicle types plotted after rescaling thickness (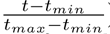) and distance (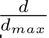) for each ossicle. The red line separates two distinct linear regimes. (N) A map of number of sides of holes represented as polygons. The number of sides decrease rapidly near the center while remaining almost constant for larger hole depth values. (O) Brightfield image of a large foot pad ossicle isolated from an adult *A. parvimensis*, by bleaching tissues. The center node is marked approximately as the midpoint of the thickest branch near the center of the plate. Foot pad is the only ossicle in the animal that continues to grow to a large size along with the increase in size of the foot. (P) Every plate like ossicle geometry displays three broken symmetries in comparison with a regular hexagonal lattice. Broken translational symmtery leads to asymmetry in the overall shape. Polygonal disorder is observed hinting any prescription of polygon shapes. And finally, radially decreasing dimensions highlight a broken radial symmetry.

To compare ossicle shapes within a single animal, we first directly quantified geometrical features including surface area, volume, and number of holes for each ossicle. To understand the principles of construction of ossicle morphologies, we investigated how number of holes varies with ossicle type and scales with material volume and surface area. We found substantial variation in the number of holes across and even within ossicle types, revealing that topological genus is not a conserved quantity during ossicle construction of same ossicle types (Figure 2D). We also observed that the volume and surface area scale directly with the number of holes, pointing to hole construction as a dominant mode of ossicle growth (Figure 2D).

To better understand the topology of a given ossicle, each 3D dataset can be skeletonized and converted into a simplified graph. This allows us to represent any ossicle structure as a lattice with distinct nodes, edges and fitted surfaces to extract subtle geometrical and topological features (Figure 2C), which can shed light on possible construction algorithms. To understand the stereotypy of geometrical prescription of a given ossicle type, we observed variations in 3D geometries of planar lattices from pillared table ossicles. We first extracted the average base plate mean curvature values for pillared table ossicles and found a global trend correlated with the local animal body curvature (Figure 2E). Given that the crystallized mineral is significantly harder (modulus *∼* 100 GPa) than the soft tissue matrix surrounding it (modulus *∼* 100 kPa) [46, 47], the curvature cannot be a result of post-growth bending but instead must be embedded during the growth history of the ossicle itself. We also observed significant variation in the size of pillared table ossicles present in different tissue niches; those in higher curvature regions, like the foot, are covered with smaller ossicles, possibly due to higher spatial crowding due to curvature (Figure S3A). This observed variation in base curvature suggests that ossicle growth conforms to the local tissue niche geometry and that variation can exist within a particular ossicle type.

Strikingly, pillared table ossicles also exhibit a range of central symmetries, that is, the number of branches emanating from the central pillared site (Figure 2F). This number ranged from three to seven in our data, though a four-pronged central symmetry was most abundant (138/147 pillared table ossicles in one animal). The absence of a strictly prescribed geometry of a given ossicle, prompted us to explore variation in its topology. We mapped the topology of each ossicle using their graph-based representation. We find that the large variations in geometry (i.e. between distinct ossicle types) directly result from variations in topology (Figure 2H). Even when we limit our scope of consideration specifically to pillared table ossicles with four-pronged central symmetry, we observe significant variation in topology with increasing node depth, measured as the shortest topological path distance from the center node (Figure 2G). It is worth noting that the center of an ossicle can often be a small segment instead of a sharply defined point, which can have significant effects on the overall topology of the network. Thus, ossicles display significant geometrical as well as topological variation across structures.

In order to shed light on this interplay between topology and geometry, we next extracted finer features of individual ossicle lattices. To understand how topology influences distance within ossicles, we compared radial and geodesic distances from the center along the ossicle surface (Figure 2I). By taking the difference between these two distance measures, it is apparent that ossicle topology enforces longer geodesic distances at distal sites. We further evaluate the local medial thickness on all points of the ossicle skeleton, which reveals a decreasing trend with geodesic distance from the center (Figure 2M), hinting at asymmetric mineral deposition across the structure during growth.

Furthermore, measured branch bifurcation angles, defined as angle between two edges (see supplementary information for details) across the structures remain almost constant around 106.52*°±* 22.34*°* for pillared tables, 107.71*° ±* 24.38*°* for other plates, and 116.45*° ±* 21.19*°* for foot pad ossicle with increasing variation in bifurcation angle with node depth (Figure 2L). This suggests that conservation of local bifurcation rules within ossicle lattices, that display increasing variance with increasing distance from the center.

By directly observing mesh geometries, we find that the ossicles are often asymmetric, extending more branches in some directions within the plane, leading to broken translational symmetry in comparison with a regular hexagonal lattice (Figure 2P). Although more prominent in front/rear plates, pillared tables also demonstrate similar asymmetries. We also find that ossicles exhibit a trend of decreasing hole areas (Figure 2J) and branch lengths (Figure 2K), constituting a broken radial symmetry. Furthermore, we find that the number of edges for each hole represented as a polygon, changes with radial distance from the center; proximal holes have more edges while distal holes have fewer (Figure 2N). This demonstrates polygonal disorder in the lattices, suggesting the absence of any regular patterns prescribing hole shapes. Moreover, across fully developed ossicles, we rarely observe edges not creating closed loops, hinting towards a tendency of the growth processes to promote hole closure. Interestingly, the foot pad plate, which grows continuously as the foot grows, displays lattice parameters that change gradually with distance/depth from the center as compared to pillared table and other plates. Thus, *A. parvimensis* ossicles are finite in size, with length and area of edges/holes decreasing towards the boundary. This variation is reduced in larger ossicles, hinting towards a coupling between terminal size of an ossicle and the growth processes generating its shape.

These observations, combined with the previously discussed geometrical and topological variations, suggest that ossicle shapes are not prescribed strictly, but rather their construction is executed through local rules, which when coupled with natural biological noise/morphogenetic cues, can result in the observed variations. To make sense of these observations, we next turned our attention towards understanding the nature of the ossicle growth process via live imaging.

### Live imaging reveals ossicle shapes as self-closing branching networks

Although geometrical and topological characteristics (Figure 2) provide insights into the construction of an ossicle, understanding emergence of these characteristics requires direct knowledge of the growth dynamics. Thus, we performed time lapse imaging of individual developing ossicles (Figure 3) in living juvenile animals at high spatial and temporal resolution. Utilizing custom designed imaging chambers and tracking microscopy, we carried out single ossicle tracking in live organisms (see methods for details) (Figure 3A) over a day. To the best of our knowledge, this is the first time live biomineral growth of individual structures has been captured in situ. Recently metamorphosed (0-20 days post larval to juvenile metamorphosis) *A. parvimensis* juveniles were observed with DIC (differential interference contrast) or brightfield microscopy under confinement at different stages of ossicle development continuously (see Methods).

**Figure 3:**
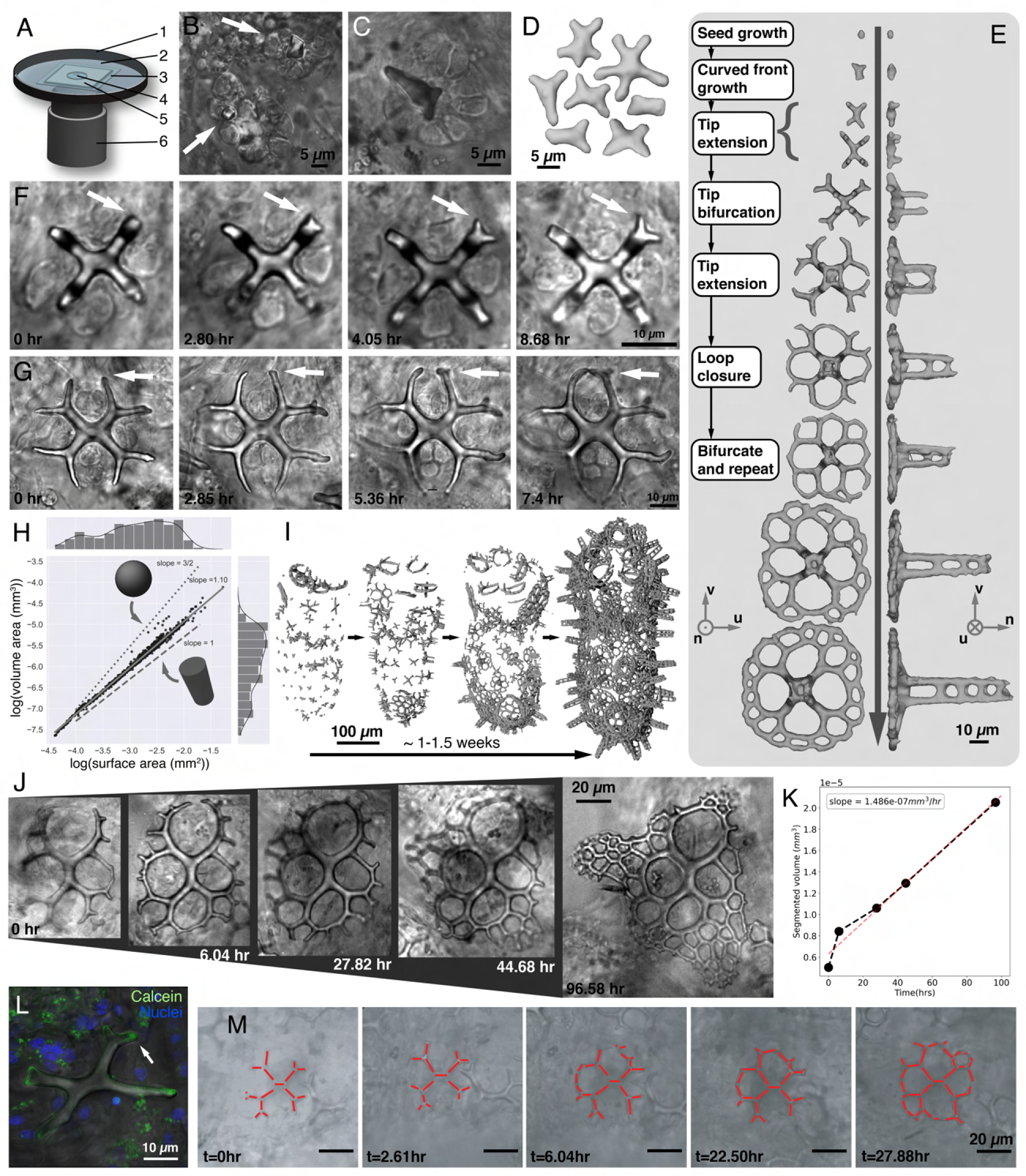
Ossicle growth happens through branching morphogenesis of biomineral. (A) Live ossicle growth imaging chamber setup. A live juvenile is slightly compressed under the weight of the glass coverslip so as to immobilize it and image the static configuration for hours. 1 - glass bottom imaging dish, 2 - fresh sea water, 3 - glass coverslip, 4 - juvenile animal, 5 - PDMS spacer, 6 - microscope objective lens. (B) DIC images of earliest ossicle seeds embedded in the sclerocyte cluster. The left one displayes rhombohedral habit associated with calcite. (C) A single ossicle seed growing curved fronts that elongate differentially to produce the first branches. (D) Extracted micro-CT data for multiple seeds showing distinct number of initial branches that can range from 3 to 7. (E) Ossicle growth happens through a series of events including seed initiation, symmetry breaking due to growth of first lobes, initial branch growth by tip extension, tip bifurcation into two new branches, tip extension, loop closure through tip fusion, and repetition of above steps. Here (u, v, n) make an orthonormal basis, with n pointing in a direction normal to the epidermis. (F) DIC imaging time series demonstrating bifurcation of of an elongating tip. Note that tip pointed by the arrow grows asynchronously ahead from the other tips. (G) DIC imaging time series demonstrating fusion between two approaching tips, pointed by the arrow. Here again, the tip closure event is seen progressing asynchronously across the four tip pairs. (H) Volume vs surface area plot on a logarithmic (base 10) scale for ossicles across different growth stages reveals a power law scaling with a slope of 1.10 which lies in between similar scalings for a sphere (1.50) and a cylinder (1.00), highlighting the near cylindrical nature of the branches. (I) Growth of ossicles across the juvenile body revealed through micro-CT data from animals at different stages of development spanning 1 1.5 weeks. Ossicle growth progresses asymmetrically throughout the body to eventually cover the entire surface. (J) Brightfield imaging time series for a large front plate ossicle growth, showing asymmetric growth of holes in multiple directions. (K) Segmented volumes for different time points are calculated through segmentation of 2D ossicle image and converting it into a union of short 3D cylindrical volumes (see supplementary information). The volume addition rate remains conserved around 1.49 10*^−^*^7^*mm*^3^*/hr* for more than 90*hrs* of growth duration. (L) Overlaid confocal and brightfield image of ossicle imaged post overnight incubation with calcein. The green signal represents additional growth happening preferentially near the tips. The dark blue label represents nuclei. (M) Bright field imgaing time series of single ossicle over the course of development reveal striking asymmetries in addition of new branches and bifurcation events, though the final structure is spatially symmetric.

We found that each ossicle is initiated as a rhombohedral habit, as previously observed in sea urchin larvae [48], resembling that of a pure calcite crystal, in association with a cluster of cells (Figure 3B). As accretive growth continues, the crystal loses its rhombohedral symmetry to grow curved fronts that differentially elongate to produce the first branches in the structure (Figure 3C, 3D). The seed can create multiple lobes ranging from two to seven in number, with four lobes being the most common, as shown in micro-CT geometries isolated for multiple seeds (Figure 3D). These seed branch symmetries lead to the distinct topologies highlighted previously in Figure 2F. These initial branches extend further and elongate, after which the tip bifurcates to produce two new branches (Figure 3F). To understand the patterns of deposition of biomineral during growth, we perform fluorescence microscopy after overnight incubation of a juvenile in calcein-labeled seawater(Figure 3L), which labels newly added calcite. We observe that the majority of fluorescence signal is localised at the tips, suggesting elongation of branches via preferential calcite deposition at the tips across the entire ossicle.

In later stages of growth, a single branch can also emerge from a recent tip fusion site, a process we refer to as ‘budding’ (Figure S2C). As secondary branches grow further, they curve and bend to meet other growing tips and fuse to form closed loops (Figure 3G). The bifurcation, budding and tip fusion events represent fundamental, universal motifs in ossicle construction. These motifs applied iteratively, with varying growth parameters, give rise to multiple layers of holes which results in the final ossicle morphology (Figure 3E, 3J). Thus, ossicle growth can be described as self-closing branching morphogenesis initiated as a small seed crystal (*∼* 1*µm*) and resulting in a topologically complex structure (*∼* 100*µm*)(Figure 3E), spanning two orders of magnitude in spatial scales.

Utilizing an extrapolated 3D geometry from snapshots during ossicle growth in Figure 3J, we evaluated the volumetric growth rate to be 1.49(*±*0.13) *×* 10*^−^*^7^*mm*^3^*/hr* (Figure 3K), which remained almost constant throughout the *∼* 96*hr* growth interval (see supplementary methods). Similarly, the surface area increase rate was calculated to be 11.36(*±*0.80) *×* 10*^−^*^05^*mm*^2^*/hr*, which also remained almost constant. This hints towards a remarkable conservation of the rate of biomineral precursor material sequestration within the short period on an individual ossicle growth. Notably, the above volumetric increase rate translates into a protein mass incorporation rate of 2.01353*pg/hr* which is two orders of magnitude smaller than an approximate upper bound on protein production rate for 6-7 mammalian cells (see supplementary information for details). Moreover, we extracted branch length extension rates from Figure 3F and G to be 0.96*µm/hr* and 0.28*µm/hr*, respectively. In general, we observe highly varying growth rates both across different animals and across different ossicles within a single animal, suggesting a dependence on local culture conditions and tissue niches respectively.

We next fixed animals at the earliest stages of juvenile growth and performed micro-CT scans, showing the maturation process (which takes 5-7 days) from early-stage seeds to fully developed ossicles at the whole-animal scale (Figure 3I). Ossicles are seeded independently across the body of the animal, with relatively uniform spacing and no direct interactions between distinct seeds. These initial seeds grow to cover almost the entire surface of the animal for the first time in its developmental history. Remarkably, each ossicle grows to a finite fixed size, after which its growth is self-terminated, thus encoding a finite length scale. As the juvenile continues to grow, new ossicles are seeded in intermediate spaces between existing ossicles but mature ossicles remain at the same size.

Next we compare ossicle volume and surface area to those of other commonly known shapes, such as a sphere and a cylinder. Extracting surface area and volume from individual ossicles across different growth stages and plotting them on a logarithmic (base 10) scale reveals a unique power law scaling with a slope of 1.10 which lies in between similar scalings for a sphere (1.50) and a cylinder (1.00), highlighting the near cylindrical nature of the branches (Figure 3H). Observing the 3D micro-CT data for similar ossicles at different stages of growth reveals that branches thicken over time, with higher thicknesses observed proximal to the initiating branch. This suggests that accretive deposition not only happens at the tips, but continues to happen across branches at a smaller rate (Figure 2M, 3E,I).

Finally, a key feature of ossicle growth process is differentially accelerated growth even amongst symmetrically placed branch tips. This promotes earlier closure of loops in specific parts of the ossicle during growth, leading to temporary, or sometimes permanent, asymmetries in the overall ossicle shape (Figure 3E, 3F, 3G, 3I, 3J, 3M). Comparing growth intermediates of pillared table ossicles, we found that each ossicle undergoes a unique trajectory of asymmetric growth (Figure 3M, S5A).

This demonstrates that any two ossicles show distinct growth trajectories (non-stereotypical) while the end results are very similar. These observations reveal a unique growth protocol which can produce symmetric objects via a branch fusion induced self-limitation on growth. It is remarkable that this is achieved in a multi-cellular cluster where no single cell has a predefined blue-print of the entire ossicle shape.

### Cell clusters create isolated privileged spaces for mineralization of ossicles

The transformation of ossicle seed into specific geometries over a short period of time motivated us to next understand the cellular context that facilitates the relevant physical processes guiding the ossicle growth trajectory. Given that a single ossicle is a large topologically complex mineral structure, it is not obvious how individual cells are able build a local coordinate system to measure its size and collectively construct its specific shape (Figure 2). Furthermore, to understand cellular control of ossicle growth, it is first critical to know the identity and architecture of the cells involved and the nature of the local ossicle growth niche. Thus, we labeled ossicles, cell membranes, and nuclei in fixed animals at different stages of growth and visualized the ossicle growth niche via fluorescence microscopy (Figure 4A,4B,4C). We identified a consistently present cell population that directly associates with ossicles throughout their development (Figure 4A, 4B, 4C). Remarkably, we observe that the entire ossicle is wrapped tightly with a thin membranous wrapping, over the course of its entire growth (Figure 4A, 4B, 4C) by a handful of cells.

**Figure 4:**
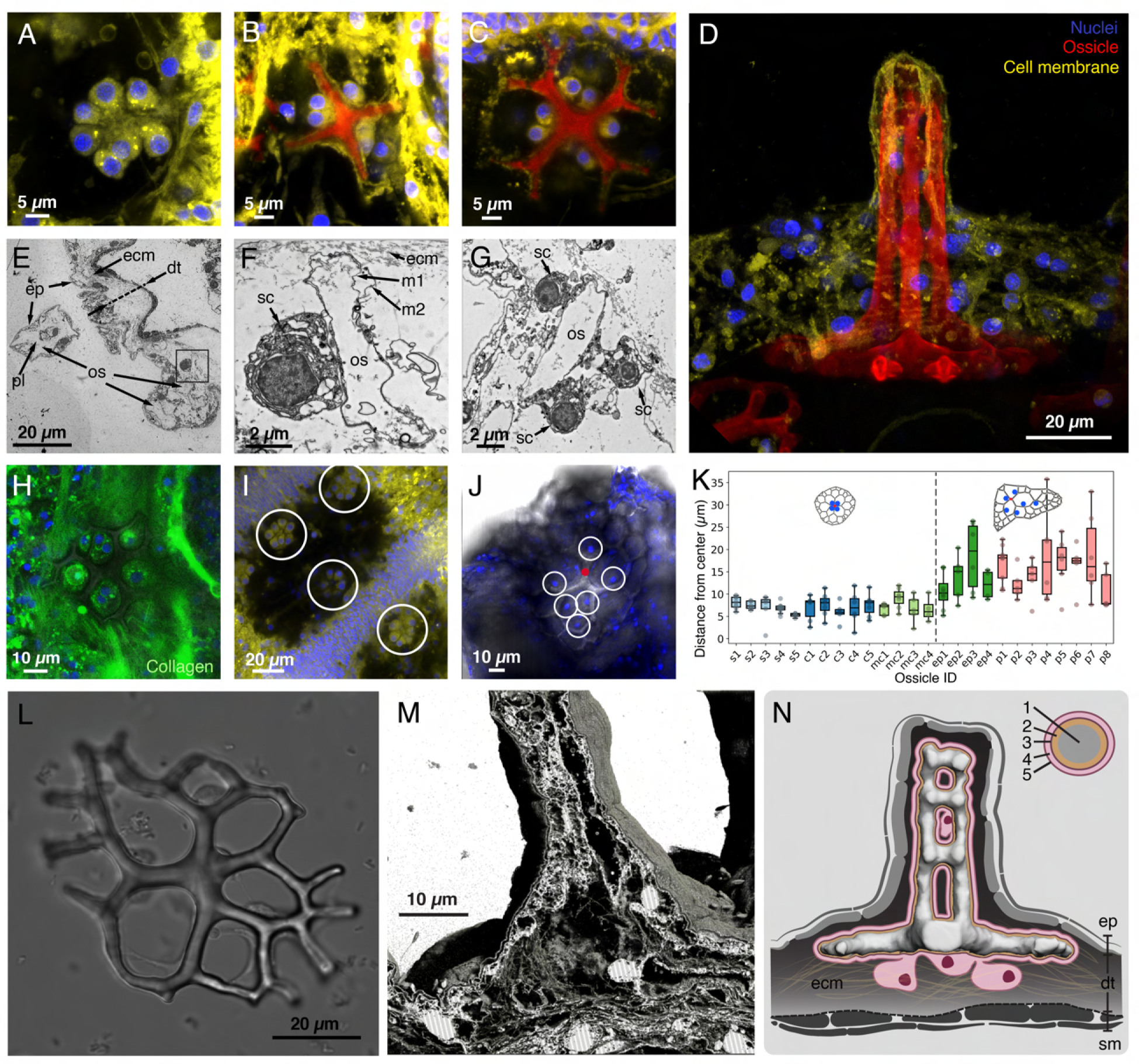
Ossicle grows within a syncytial confinement created by a distinct cluster of sclerocyte cells. (A-C) Confocal scan images of sclerocyte clusters from distinct ossicles mapping the growth progression of a pillared table ossicle (Yellow – Cell Membrane, Blue – Nuclei, Red – Calcein labeled ossicle). Ossicle growth begins in a 4 10 sclerocyte cell cluster. As the ossicle grows, the cells maintain a continuous cytoplasmic wrap throughout the surface of the ossicle, even during binary splitting and tip fusion. (D) A maximum intensity projection of confocal z-stack of side view of pillared table ossicle from a fixed juvenile. It shows clear absence of any tissue right below the ossicle, and the vertical pillar protruding out, while being wrapped by the epidermis. (E-G) TEM imaging of 80 nm thick sections of a resin-mounted whole juvenile *A. parvimensis* at different scales of observation. In E, the entire dermal tissue (dt) section is visible along with a slice through of a pillar (pl) of a table ossicle. The clear empty regions represent cross sections of ossicles (os), which got dissolved during the fixation and embedding process. The dense filamentous features represent collagen dense extra-cellular matrix (ECM). The epidermis (ep) maintains and acts as the barrier to the external environment.

Previously, these ossicle-associated cells have been characterized as sclerocytes and likened to the larval sea urchin primary mesenchyme cell (PMC) population, which is known to drive mineralization through sequestration of salts from the sea water and production of organic matrix molecules [41, 42, 49]. However, the spatial arrangement of sclerocytes along the ossicle and its role in regulating ossicle geometry has remained elusive. Furthermore, the underlying cellular processes driving growth of specific ossicle geometry in holothurians remain elusive.

To determine the exact architecture of the sclerocyte cell clusters, we performed TEM imaging of fixed tissue sections, which revealed a remarkable continuous wrapping of the ossicle by a double-membrane bound ultra-thin cytoplasmic sheath (as thin as 40 nm in some regions) (Figure 4E,F,G). As we traversed around the surface of the ossicle within and across different TEM sections, no cell-cell junctions could be observed, suggesting continuity of cytoplasm across cells, which is a hallmark of a syncytial architecture as previously suggested [41, 42]. Syncytial confinement enables the creation of a protected environment for biomineral growth, which cells can then access and regulate, similar to other ‘privileged spaces’ in biomineralizing systems [2]. In order to rigorously test the existence of a syncytium, we dissociated dermal tissue with collagenase and calcium-free seawater and found that sclerocyte cell bodies and membrane wrapping remained attached to the ossicle post-dissociation (see Methods, Figure 4L). Thus, we conclude that this is a true syncytial architecture.

We next considered the broader tissue context in which ossicles grow. In the later parts of doliolaria stage (Figure S1A), multiple sclerocyte clusters composed of about two to ten cells (six being most common) appear within the dermal tissue across the entire body of the animal, initiating distinct ossicle growth sites (Figure 4I). Simultaneous growth within multiple sclerocyte clusters results in a discrete distributed skeletal structure with hard ossicles embedded in a soft catch connective tissue made of collagen [50]. Ossicle-sclerocyte syncytia are located within the body wall, sandwiched between epidermis on one side and the collagen matrix on the other, in a disc-like configuration (Figure 4N). The seed crystal begins to grow within the syncytium, positioned slightly above the cell bodies (Supplementary Video SVX). Further planar growth happens right below the epidermis and above the collagen bed while the vertical pillar grows perpendicularly as the epidermis conforms to its new shape (Figure 4D,M). We also visualized collagen and non-specific extra-cellular matrix (ECM) organization via DIC and fluorescence microscopy (see methods) and did not observe any patterns neighboring growing ossicles that resemble fenestrated ossicle geometry (Figure 4H, S4B,C). We did observe direct connections between the ossicle and ECM fiber bundles which potentially maintain the relative positions of the ossicles within the animal as it contracts during movement and defense manoeuvres. This rules out any role of ECM-based patterning to directly guide or template ossicle growth, but indirect effects due to arrangement of ECM fibers near growing tips cannot be ruled out. Moreover, morphogenetic cues from the local tissue niche, primarily epidermis and other neighboring sclerocyte clusters, may guide the development of the ossicle, which has not been explored here.

Given that sclerocyte cell bodies are likely sites of material sequestration and directors of transport processes, their spatial positioning along a developing ossicle could greatly influence the growth process. Thus, we next investigated the spatial organization of these cell bodies during ossicle growth. Confocal imaging of live and fixed tissues at different stages of growth revealed the spatial positioning of syncytial cell bodies along the ossicle (Figure 4J, 4K). Interestingly, we found that cell bodies remain localized near the center of pillared table ossicles over the course of their entire growth (Figure 4K), rather than being motile or localizing to the growth sites at the tips. This can also be observed in time series presented in Figure 3F and 3G. On the other hand, cell bodies associated with flat front plates localize distally from the ossicle center. The sclerocytes need to support vertical growth in pillared table ossicles and hence a central position could give them access to support growth in both planar and vertical directions. In contrast, an absence of vertical growth in front plates could allow the cells to spread out planarly and support larger planar growth. This hints towards the role of cell body positions in regulating growth through local distribution of resources and thereby determining ossicle shape.

A cellular syncytium is an effective strategy for maintaining a protected zone for biomineral growth [2]. This approach allows cellular control over biomineralization chemistry and provides access to the growing ossicle for continuous growth, but it also has consequences for cell geometry and architecture. Namely, the high curvature regions of the ossicle (for example, growing tips) must introduce regions of high membrane curvature into the cell, which could directly affect ossicle shape sensing and material transport processes. Put differently, sclerocyte syncytium becomes the shape of the ossicle it is building and ossicle growth continuously responds to changes in the syncytium shape, creating an interesting opportunity for feedback between ossicle morphology and cellular control of growth.

In F, a single sclerocyte (sc) cell wraps the ossicle section with a continuous double membrane-bound cytoplasmic sheath. The thickness of the sheath varies along the surface of the ossicle but can reach the smallest dimensions of up to 40 nm. m1 and m2 represent the inner and outer membranes respectively. In G, three sclerocytes join to continuously wrap their cytoplasm around the ossicle section. No membrane break is visible in the cytoplasmic path, which corroborates previous observations [41, 42] that sclerocytes form a syncytium. Thus, the ossicle grows in a membrane-bound vesicle-like region within the syncytium made by the sclerocytes. (H) Confocal scan section of an ossicle (dark halo) embedded in the ECM. Collagen rich fibers (Green) penetrate the local neighborhood of the ossicle, but do not assume any pre-patterns that the ossicle could use as templates for growth. This suggests that the ossicle growth happens by continuous modification of the nearby ECM and is only locally prescribed by the sclerocyte cells and their neighborhoods, rather than a global prescription by the matrix. (I) Multiple sclerocyte clusters (in circular rings) develop simultaneously across the surface of the animal in the late doliolaria stage. The clusters are separated from each other enough to prevent overlapping growth. The dark blue patch of nuclei represents the ciliary band in the larvae. (J) The cell body distribution in the front plate ossicle is distinct from the table ossicles. The presence of the central vertical pillar is correlated with the more centrally located cell bodies in the table ossicle. In comparison, cell bodies for front plate ossicles shown with blue nuclei in the image disperse more from the center supporting the enlarged planar growth of the ossicle. This suggests a need for proximity dependent availability of cellular resources to contribute to growth fronts in both planar and vertical directions. (K) Box plot showing planarly projected relative positions of sclerocytes from ossicle center extracted using brightfield and confocal images of 26 ossicles. For pillared table ossicles, the cells remain localised near the center to support both vertical and planar growth. For front plates, cells disperse much further away from the center. Distinct sample ids refer to: s - seed, c - cross(pillared table), mc - mid-closure(near fusion, pillared table), ep - early plate (front plate), p - plate (front plate). Distinct colors represent these distinct ids. (L) DIC image of an ossicle isolated from the rest of the tissues using a collagenase treatment. It displays topologically linked sclerocyte cells that form a syncytial wrap around the ossicle. (M) 3D segmented rendering of SBF-SEM dataset from table ossicle of a resin embedded juvenile. It shows numerous collagen fibers and membranous projections covering empty space representing dissolved ossicle. (N) Cartoon representing a longitudinal view of a table ossicle, with sclerocytes and local tissue niche. The inset represents the cross section of an ossicle stem where 1 – ossicle, 2 – ossicle - membrane space, 3 – inner membrane, 4 – cytoplasm, 5 – outer membrane. The yellow streaks resemble collagen rich ECM. ep – epidermal tissue, dt – dermal tissue, sm – smooth muscle.

### Active cellular processes enable ossicle construction

Long term observation of syncytial sclerocytes reveals a near static localization during ossicle development. Since the cell body itself is not crawling on the surface of the ossicle, how does an individual sclerocyte control the location of deposition of minerals and organic matrix molecules. This challenge motivates us to investigate cellular transport processes in sclerocytes enabling material transport. The non-uniform accretive growth enabling branching morphogenesis of ossicle geometry requires preferential material deposition at growing tips. Furthermore, given that sclerocyte cell bodies, the assumed sites of material sequestration, are positioned far from growing tips (Figure 4K), they must be able to solve the challenge of long-range directed transport to the constantly evolving deposition sites. Next we perform live imaging of developing ossicles and scleroyctes at high temporal resolution along with fixed imaging of intracellular architecture.

Live confocal imaging of the ossicle growth niche after a short (*∼* 2 hrs) calcein pulse reveals *∼* 0.1 *–* 1 *µ*m puncta of calcein decorating the ossicle and specifically localized within the cytoplasmic sheath (Figure 5A). Similar structures were observed with membrane labeling, suggesting their identity as membrane-bound vesicles (Figure S8A). For a more detailed view of these membrane-bound vesicles, we performed TEM of ossicle sections and observed similarly sized vesicles localized within the sclerocyte cytoplasmic sheath (Figure 5B). These observations suggest that sclerocytes transport precursor materials for ossicle growth in small vesicles localized within the cytoplasmic sheath and therefore restricted to the surface of the growing ossicle. Vesicle transport of calcium carbonate has been implicated in larval sea urchin spiculogenesis [49, 34]. In order to release mineral precursors into the ossicle growth compartment, these vesicles must fuse with the inner membrane (see Figure 4M), which has been shown to occur in larval sea urchin [34]. Furthermore, it has been demonstrated in several systems that the released amorphous precursor material must then undergo a physical transformation into calcite, and a similar process is likely to occur here [48].

**Figure 5:**
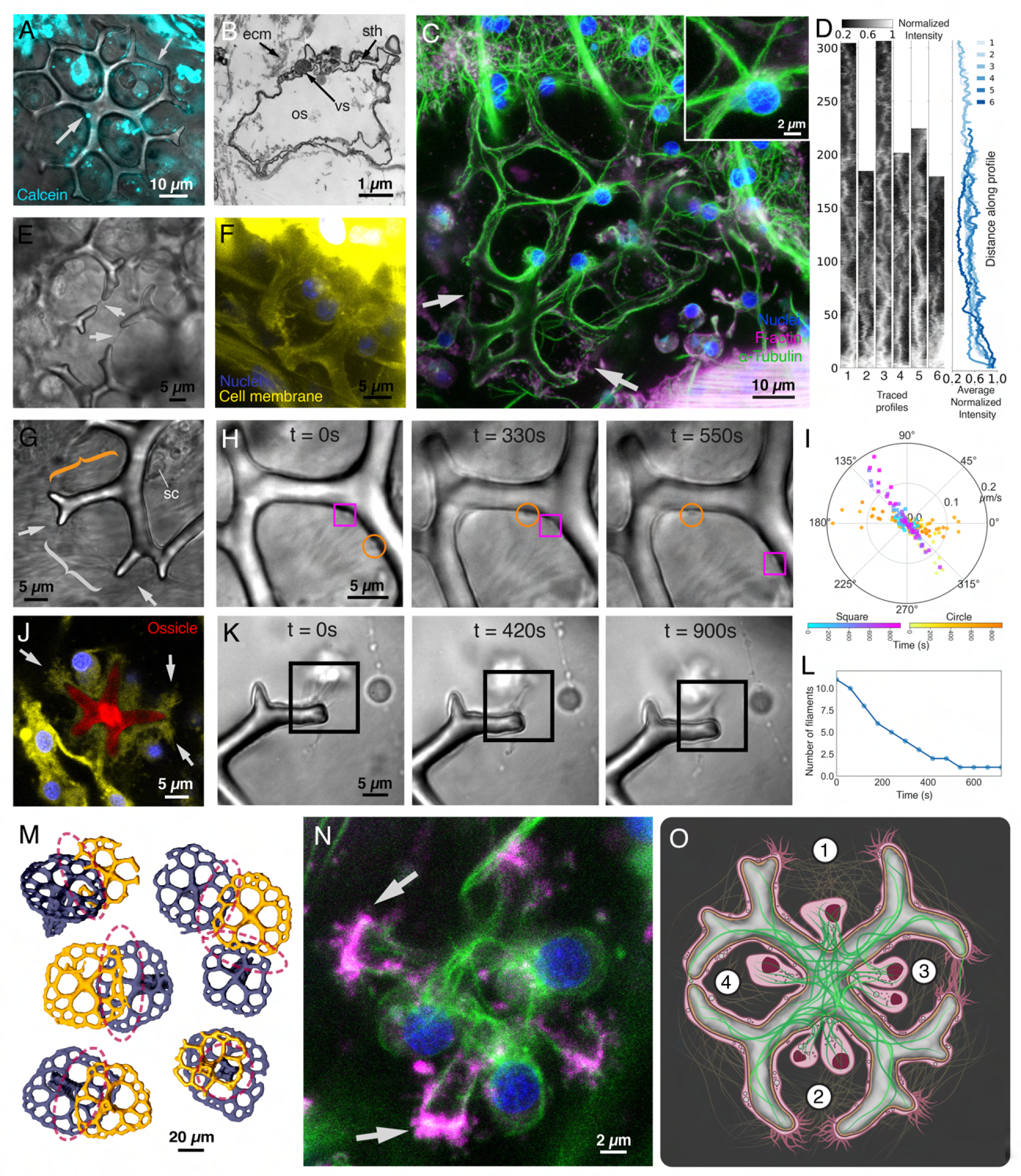
Filopodia-like activity on growing tips, and transport restricted to the surface enable ossicle morphogenesis. (A) Confocal imaging of motile vesicles (Cyan) labelled with calcein located within the syncytium. The vesicles demonstrate a non-uniform size distribution and localization within the syncytium. (B) Circular vesicle-like object appearing to merge with the inner membrane, in the cross-section TEM of an ossicle branch. The vesicles must fuse their membranes this way in order to release their contents into the ossicle space. vs-vesicle. (C, N) Maximum intensity projections of confocal scan z-stacks of ossicles from fixed animals showing actin filaments (Magenta), microtubules (Green) and nuclei (Blue) (See supp. video X,Y for traversal of the z stacks). N shows an early stage ossicle with high density of actin filaments at the growing tips along with microtubules emerging from the sclerocyte cell bodies, wrapping the central region and spreading out to the tips. C shows a larger front plate ossicle with long microtubule bundles emerging from the cell bodies and spreading far out to the furthest tips, covering the entire ossicle. The microtubule density is highest near the cell bodies but diminishes as we go further towards the tips. The arrows point towards instances of microtubule and actin filament presence across membrane threads between approaching tips. (D) 15 pixel wide paths traced along the microtubule bundles to capture the decreasing intensity with distance from the cell bodies. The right panel shows average intensity along each 15 pixel row plotted against the distance from cell body. (E) Membrane threads between approaching tip pairs in two nearby ossicles. The tips ensure fusion with nearby tips from the same ossicle, irrespective of the close presence of tips from other ossicles. (F) Confocal section of live membrane (Yellow) and nuclei (Blue) labeled ossicles showing membrane threads between approaching tips. (G) Nearby branch tips with filopodia-like active projections (arrows) that are reaching out to connect and form a membrane thread (white brace). Surface-restricted cytoplasmic activity is visible in the labeled (orange brace) branch (see supplementary video X) along with the participating sclerocyte (sc). (H) DIC imaging time series of two boluses, with similar sizes as vesicles moving past each other as they glide with punctuations along the surface within the cytoplasmic sheath. (I) Polar plot showing velocity distribution for both the tracked boluses. (J) Confocal section of an early stage ossicle within a fixed juvenile, displaying membranous projections at the tips. The right tip displays a membrane projection bifurcation possibly preceding a mineral bifurcation growth. Yellow - membrane, Blue - nuclei, Red - ossicle. (K) DIC imaging of a tip of an ossicle freshly separated from the rest of the tissue using collagenase treatment. The tip displays multiple filopodia-like membranous projections that are otherwise obscure in ossicles imaged *in vivo*. The time series depicts merging of multiple projections into a single one, gaining similar morphology to membrane threads between approaching tips. (L) Plot displaying decreasing number of filaments with time, visually identified in series K. (M) Self-recognition of ossicles leads to asymmetric growth, with growth restricted in regions adjacent to other ossicles nearby. The ossicles labeled in yellow represented ossicles with affected growth due to proximity to the purple ossicles. (O) Schematic highlighting cellular physiology and mechanistic steps involved in the ossicle growth, namely 1. branch extension and tip search, 2. tip approach and elongation, 3. membrane thread formation with cytoskeleton penetration, and 4. membrane fusion and loop closure by the mineralized phase. The green lines represent microtubules and the yellow lines represent collagenous substrate.

Next, we investigated how vesicles are transported around the complex geometry of the ossicle within the cytoplasmic sheath. To address this question, we performed high magnification (*×*60 *×* 1.5) live DIC imaging of immobilized juveniles, which reveals active shuttling of cytoplasmic cargo (largest sizes up to *∼* 1 *−* 2 *µ*m) restricted to the surface of the ossicle (Figure 5 G,H, Supplementary Video SVX). Based on the size and localization of this cargo, we presume that these are the same vesicles observed in Figure 5A,B.

We extracted velocities of two distinct boluses (Figure 5H), and observed velocities of punctuated strides ranging in the order 0.01 *−* 0.2 *µ*m*/*s (Figure 5I). We additionally observe smaller puncta moving at much higher velocities [see movie XYZ]. Notably, this finding is consistent with previous observations in neuronal transport with rapid unidirectional movement of submicroscopic vesicles [51, 52, 53].

Traditionally, spatiotemporal reorganization and transport of vesicle cargo and organelles is often achieved through cytoskeletal structures and motor proteins. This has been extensively highlighted in the context of trafficking, secretion and signaling [54, 55], especially in neurons [56]. Moreover, distinct cell types in metazoans are known to utilize unique arrangements of oriented cytoskeletal arrays to support specific cellular functions [57]. Given the unique multi-holed topology of the cellular syncytium around growing ossicles, we next explored its cytoskeletal architecture. Through immunolabeling (Figure 5C, 5D), we discovered an extensive network of microtubules radiating from the sclerocyte cell bodies that span the entire surface of branched ossicles. The microtubule networks are present in early stage ossicles (Figure 5N), in intermediate growth stages (Figure S9), as well as in well-developed large ossicles (Figure 5D,S9). We find that the microtubules are arranged in bundles, oriented longitudinally along the ossicle branches. These bundles appear to spread and thin out with distance from the cell bodies, with individual microtubules dispersing and extending to span all the branches. We quantify this in Figure 5D as decreasing relative intensity of microtubule bundles with increasing distance from cell bodies. Such an arrangement of microtubules is conducive to 1D transport of vesicle cargos (Figure 5H), akin to their role in axonal transport [58] in a topologically complex cell. The presence of continuous microtubule network across the ossicle can enable the transport of packaged precursor amorphous mineral and matrix proteins from the cell bodies to the growing tips. The lossy nature of cargo transport along microtubules, stemming from spontaneous or competitive detachment of motor proteins, termination of filament tracks, and organelle congestion [59, 60, 61, 55] also enables strengthening and accretive growth increasing the thickness of existing branches. Thus, the oldest segments of branches would grow to become the thickest due to their presence along the paths transporting material to the tips, resulting in a decreasing thickness profile from the center to the tips, ass shown previously (Figure 2M).

To further elucidate the cytoskeletal organization of sclerocytes, we investigate the spatial organization of f-actin via phalloidin staining. F-actin is visible across the the cytoplasmic sheath and surprisingly, high density of filopodia-like active projections are present at the growing tips (Figure 5N, S8). Moreover, we observed thin thread-like projections connecting distant regions of branches within a single hole, across the ossicle (Figure S8A,B). We confirmed these observations through direct observation of membranous activity at the growing tips using DIC microscopy. The filopodia-like membrane projections (length around 5 *µ*m) radiate outward from the growing tips into the surrounding matrix and display activity through membrane fluctuations occurring at a time scale of tens of seconds (Figure 5G, Supplementary Video SVX). We did not observe such activity universally across all branch tips, suggesting cellular agency in tuning this activity. For higher contrast imaging, we partially isolated one of the developing front plate ossicles with a tip displaying filopodial extensions using transient collagenase treatment. The higher contrast achieved due to separation from the background field of collagen fibers revealed multiple fluctuating filopodial projections emanating from the branch tip, slowly coming together over the course of *∼*15 minutes to give rise to a thin bundle (Figure 5K,5L). Moreover, membrane labeling reveals similar filopodial bundles at growing tips as shown in Figure 5J. One such bundle was captured exhibiting a bifurcated geometry, hinting towards the possible role of these active projections in guiding bifurcation events. Such actin-based, filopodia-guided bifurcation of membrane extension could be further stabilized by microtubule growth, as has been demonstrated in cytoskeleton driven collateral branching in neurons [62].

In some cases, we observed a thin thread-like projection connecting two neighboring growing tips (Figure 5E). Such projections show membrane labeling (Figure 5F), as well as cytoskeletal labeling for both microtubules and actin (Figure 5C). A single microtubule usually passes through the membrane-bound thread, connecting the two tips, along with actin filaments organized around it (Figure 5C, white arrows). The presence of such projections is also visible in time series in Figure 3G, between the approaching tips. The absence of such threads in other pairs of neighboring tips that are further separated, suggests their dynamic formation during ossicle growth in a distance-dependent manner. Such pairing between the tips appears to enable asymmetric localization of mineral precursors and curved growth of tips towards one another. It is possible that the filopodial extensions observed in Figure 5N, if present in two neighboring tips, could interact to form these threads, though these dynamics have not been investigated here. Thus, we hypothesize that tips with active filopodia-like membranous projections search their local neighborhoods and pair up with a nearby tip by extending projections and fusing them to create a membrane thread that further supports the mineral growth and fusion of participating tips.

Surprisingly, despite the extensive cellular activity discussed here and high-density placement of ossicles in the body wall, no cases of inter-ossicle merging were observed in our light microscopy observations of *>* 20 animals with hundreds of individual ossicles per animal. This observation is also consistent with all the micro-CT scans of individual ossicles from *>* 5 animals. Even in cases where adjacent ossicles exhibited overlap or linking (Figure 5M), no tip merging was observed. This suggests a strong “self-recognition” capability within every sclerocyte syncytium, which could be mediated through molecular sensing, developmental gradients, or mechanical interactions with the nearby ECM. The exact mechanism for this remains unknown. Beyond self-recognition, nearby ossicles also inhibit the free extension of growing tips, causing reduced or altered growth in direction where a neighbor is present (Figure 5M). This self-recognition and neighbor inhibition is crucial in context of sea, since they exhibit discrete distributed skeleton without formation of fused structures, as observed in other echinoderms.

In summary, ossicles are topologically complex branched mineral structures wrapped by syncytial sclerocytes which utilize a microtubule network to transport vesicle-bound material to mineral growth sites. Furthermore, actin driven filopodial dynamics at growth tips guide and regulate fusion of branches, thus guiding morphogenesis. The critical role of top-down processes and cellular physiology in ossicle morphogenesis leads us to hypothesize that ossicle shape is governed by not only bottom-up processes such as mineralization at growth sites, but also top-down cellular processes that guide material to specific sites on the surface of the ossicle. The continuous maintenance and replenishment of resources [63] governed by sclerocyte cells allow us to explore a top-down control of finite ossicle geometry. To test this hypothesis, next we pursue a material transport model to explore how constraints on cellular resources and calcium carbonate sequestration can affect emergent ossicle geometry.

### Resource-constrained growth of a self-evolving branching network

Motivated by the interplay between cellular constraints and branching morphogenesis, we decided to simulate material transport coupled growth of a self-evolving discrete branching network. For a tractable model, we simplify the ossicle shape dependent organization and transport over microtubules within the sclerocyte syncytium to a graph network (where edges still have finite thickness enabling us to represent an ossicle). We construct a minimal model that allows us to capture the processes of directed transport-enabled tip growth, bifurcation of tips, active tip guidance and loop closure progressing under a resource constraint. Our model sufficiently captures the transport dependent feedback between the evolving syncytium and ossicle geometries and allows us to highlight the universality underlying the observed ossicle shapes.

We first write a continuum model of linear tip growth using a simple 1D model aligned along the X axis. The idea is to build intution over the process of lossy directed transport over an existing structure (modeling microtubule directed transport of vesicles). Let’s assume that mass is being injected at a point, say the origin in this 1D model, at a constant rate 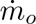 (measured experimentally in Figure 3K). Thus mass (captured in discrete vesicles but modeled here as continous transport problem) gets directionally transported towards the tip while a fraction of it gets dropped due to leaky transport (fusion of vesicles to the outer membranes) along the path length. This simple process mimics the lossy nature of vesicle transport over microtubule architecture. Whatever mass reaches the branch tip (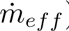) contributes to its linear growth while any mass deposited along the way contributes to the increase in thickness of the structure. The decrease in mass rate due to fractional dropping can then be locally written as 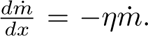 Here *η* is the fraction of total mass dropped when it traverses over a unit distance. An alternative extreme of this could be constructed using an assumption that a constant amount of mass, 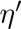, gets dropped every unit distance, in which case the mass rate change equation becomes 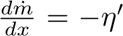. The time dependent shape profile of the growing branch can then be evaluated to be 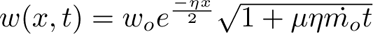 for the former case and 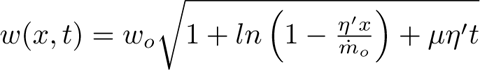 for the latter respectively. Here 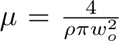 is a physical parameter that decides the branch length increase due to a unit mass accumulation at the tip, *ρ* is the final density of material post incorporation within the ossicle, and *w_o_* is the minimal initial width as the tip grows. We can observe that the width at any position grows proportionally to 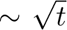 and the profile is dependent on the mass injection rate as well as the mass drop/loss fraction. This simple 1D model provides intutive scaling for both growth of the tips and thickening of the branch. Next, we extend this simple abstraction over complex graph networks/topologies resulting from bifurcations and fusion of tips.

In order to capture the growth dynamics in ossicle geometries, we numerically solve for the material transport and local growth over discrete branched networks composed of nodes and edges. Transport on this graph network allows for the graph to grow as a function of time, while the connected nature of this graph directly regulates transport via a growing topology. We initialize each simulation with a graph representing the initial seed displaying a broken symmetry with 4 lobes growing within a single sclerocyte cell cluster. We assume that the precursor material produced by each cell is injected at the rate 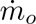 at the location marked by the cell body (Figure 4A,B,C) on the existing graph (Figure 6A). Mass is transported by vessicles along the existing graph away from the cell node and towards the tips. Bio-mineral precursor sequestration, packaging into vesicles and transport intracellularly in the synctium are all resource-limited active processes performed by the sclerocyte cells which together determine the total injected mass rate 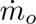. We incorporate these resource constraints on multiple active cellular processes using a single parameter in our simulations. The resulting mass flux arriving at any given node in the graph are then evaluated by solving for the loss resulting from the fractional drop at each edge (similar to the 1D model) which contributes to the growth of their width, *w*, at each time step. The total mass contribution from multiple cells reaching at the growing tips during each time step, 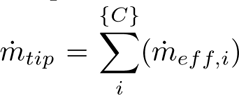 is then used to evaluate tip growth within that time step 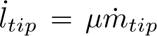. The growth of the tip is processed through addition of new nodes at fixed separation and saving any remaining mass at the tip for the next time step. We also incorporate the tip attraction and splitting events we notice in experimental framework via a simple set of tip rules. Tips continue to grow until 1) they touch a non-growing structure, in which case they fuse and terminate; 2) they touch another growing tip in which case they either fuse and terminate or bud a single tip post-fusion; or 3) they undergo binary splitting into two growing tips. The directionality of transport over each edge of the network is enforced to be the direction along which the edge has grown during the evolution of the network, mimicking the directional laying of microtubule bundles during growth. A bifurcation or fusion event thus enforces equal distribution among daughter branches or merging of mass fluxes respectively. In order to simplify the simulations and focus on emergent geometrical features of a growing graph network, we implement a sychronous tip bifurcation rule at a mean angle of 108 deg (Figure 2L). Although asynchronous bifurcation events do occur at irregular intervals, as imaged in live time-series ossicle growth experiments (Figure 2G, J, L), a synchronous bracnhing model still captures the overall geometry/topology of the growing network.

**Figure 6:**
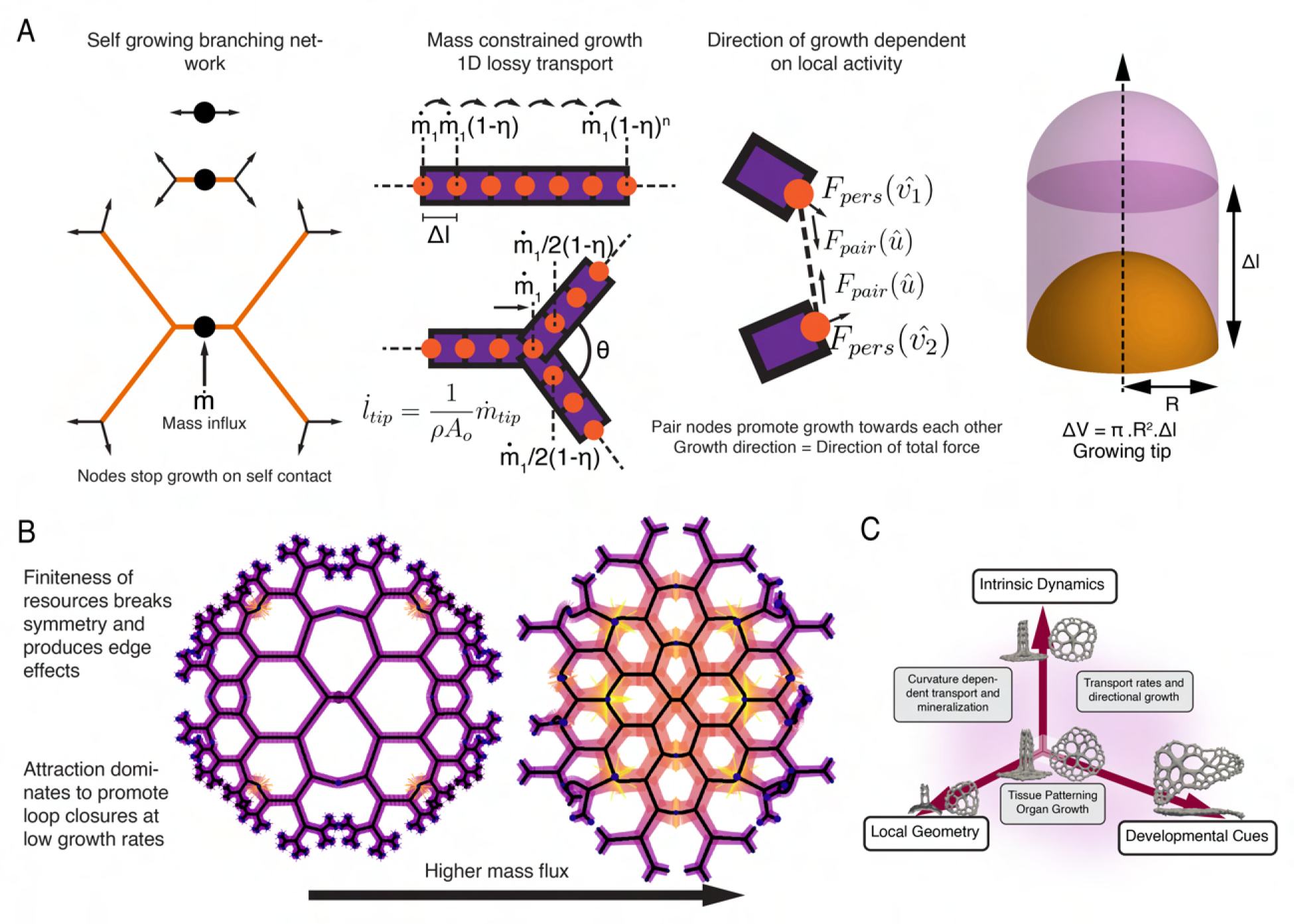
A resource constrained self evolving branching network produces ossicle-like geometries. A network is initiated at a single node that has bifurcated into four branches creating the initial asymmetric structure. Mass is assumed to be injected at the central node representing cell position, at a constant rate, which gets transported to the tips while getting dropped on the way at a constant fraction. Tip growth happens in directions prescribed by the net force acting on them, which takes into account all the local and global information sources. The tip is assumed to extend with a minimal constant thickness which later thicken due to lateral deposition. Branches are assumed to bifurcate at constant intervals for simplification. The branches fuse and terminate which leads to formation of closed loops. This process results in a structure similar to pillared table ossicle geometry, displaying decreasing hole and branch dimensions radially. This rate of variation of dimensions can be altered by tuning the mass influx rate. A higher mass influx rate allows conservation of effective branch extension rates between bifurcation events, resulting in geometries similar to larger foot pad ossicle. Thus, intrinsic dynamics of the ossicle-sclerocyte com-plex, the prescription of space by the local geometry and external developmental cues from neighboring tissues (not studied here) couple together to generate the space of morphologies observed in the animal.

In order to capture the observed curved growth of tip merging events, we maintain a direction vector *θ* at each growing tip that dictates the direction in which new nodes are added in a given time step. This direction is dictated by the filopodial activity-enabled active tip guidance which is potentially dependent on the cumulative effect of multiple morphogenetic cues being sensed by the filopodia. To simulate the dynamics of active tip guidance we utilize an abstract dimensionless forces acting on the tips whose resultant vector determines the directional vector *θ* for growth. Firstly, we assume a persistence force, *F_pers_*, acting on each tip that ensures a tip to continue to grow in a direction defined by the immediate bifurcation from which it was born, capturing a local control of ossicle growth direction. This assumption is based on the observation of near straight growth in tips that are far away from any other tips and can be justified due to the energetic costs involved in bending membrane and microtubule filaments during growth. Secondly, as shown in live imaging of highly dynamic membrane filopodial networks at tips (Figure 5K, N), we assume that growing tips interact in a distance-dependent manner. This represents the active search observed in nearby non-sister tips (tips that have not split from the same parent tip) to find a neighbor and form a closed loop. We further assume that each growing tip maintains an active search radius defined by *R_o_*. Any other non-sister growing tip in this active search region can also be used to pair up and make a membrane bridge. Once a bridge is formed, the participating tips do not search or make any further bridges. To incorporate this highly dynamic tip-search algorithm (akin to growth cones in neurons), we model a distance independent directing force, *F_pair_*(*u*^), on the tip oriented along the relative position unit vector towards the pair tip, *u*^. The two tips thus keep moving towards each other until they merge and growth is terminated. In summary, in the absence of any interactions, each tip persistently continues to grow in a direction defined by the immediate bifurcation from which it was born. Whereas any interaction with the neighboring active tips changes the net angle of a growing tip.

The effective halving of available mass rate for tip growth following each bifurcation results in progressively decreasing tip extension rates as the structure grows further away from the center (also seen in the experiments). This results in shorter branch lengths under a constant splitting interval timer. Tuning the available mass flux near growing tips due to addition of more cells, or due to constant shifting of the position of the cell bodies over longer time scales, results in a near uniform field of hole sizes and branch length, mimicking a feature seen in foot pad ossicle. Variations of baseline graph network topologies are possible via processes that would interact with growing cellular fliopodial tips, further motivating assymetric growth. Thus, cytoskeletal-mediated directional transport can give rise to asymmetric growth in the shapes similar to those observed in front plates (Figure 6B).

Although simplified in nature, the above simulation framework allow us to develop a deeper understanding of how multiple cells confined to a specific geometry of the ossicle as a syntium, can enable a top-down emergent construction of biomineral lattice where it’s corresponding graph structure controls transport and in a close feedback - the directed transport controls the graph network. As an example, the exact locations of cell bodies in our model drives local mass injection within the growing branching network, resulting in a relative longer edges in the neighborhood of the cell body and a field of progressively decreasing edge lengths further away, as observed in statistics extracted from micro-CT data (Figure 2). Moreover, overlapping material flux from multiple cells leads to an emergent edge length across larger length scales, as seen in large scale foot pad ossicles.

## Discussion

We have introduced juvenile *Apostichopus parvimensis* as a model system for studying morphological programming of biomineralized ossicles in sea cucumber. Since hundreds of ossicles grow simultaneously throughout the body of a juvenile organism, the presented experimental system allows us to ask novel questions about ‘cellular masonry’ or how cells collectively construct complex biomineral shapes. We utilized a range of experimental and theoretical techniques to study the fundamental principles of holothurian ossicle development establishing an ideal system, which can be further extended in future studies. For example, comparative micro-CT scans across species would will shed light on the universality of ossicle construction rules presented here. Ossicles present a unique case for geometry-coupled transport-induced growth, observed ubiquitously across multicellular contexts. Because prescription of form does not happen directly at the length scale of the entire ossicle, information is read locally across the growing structure by the cellular processes, and overall geometry emerges out of these local interactions. This results in the natural noise in the system, due to locally varying execution of growth processes. Scanning a large number of ossicles would enable rigorous quantification of error rates associated with morphogenetic processes that pattern individual ossicles. Finally, local perturbation of individual ossicle growth, through chemical (cytoskeletal drugs, quenchers, pH) or physical (temperature, compression) disruptions, while using other ossicles in the same animal as a control, would allow us to build deeper understanding of the molecular processes at interplay between biological and physical layers of complexity presented here. By comprehensively mapping ossicle growth, shape, and form across the entire animal, we present the first 3D morphological atlas of sea cucmber ossicles mapped to specific parts of the organism’s body plan. As highlighted in Figure 6C, ossicle shapes can be generated through intrinsic cellular dynamics coupled with local geometry and developmental cues to guide ossicle tip growth, with cell position dispersion regulating variations across ossicle types.

In context of biomineralization studied across the tree of life, mineral biomolecular content has often been suggested to play a key role in guiding growth [64, 65] as well directly prescribing complex shapes the mineral structure can take[66]. However, how extremely local processes depositing organic material at local scale can program global ossicle shape remains poorly understood. Here we present a top-down cellular control of transport processes as another guiding principle for shape programming in biomineralization.

The accretive growth of biomineral geometry presented here is fundamentally distinct from other branching processes observed in nature in three key ways. Firstly, branching networks found in leaves, trees, capillary networks, river source or delta networks, and slime molds are shaped by the flows that they support [6, 67], which are proportional to *d*^2^, where *d* is the thickness of the branch. In contrast, material flows during ossicle lattice formation via surface restricted transport, can only scale with a factor of *d*, enforcing a geometrical restriction on the growth dynamics. Secondly, branching networks within organs such as lungs, salivary glands, and kidneys involve non-uniform surface or bulk tissue growth induced by morphogenetic coupling between the tissues and their local environments [68]. In these cases, material injection and bifurcations happen through tissue scale cell division-induced growth, whereas ossicle branching relies on cell scale activity for the same (Figure 4-6). Thirdly, solidification/precipitation induced growth of dendrites in melt pools/supersaturated salt solutions relies on material accretion from the entire bulk neighborhood, thereby preventing the merging of branches due to depletion of salt or metal contents from the media between two approaching branches [7]. Due to the presence of a surface restricted environment for the growth of mineral phase in ossicles, the approaching tips do not deplete concentrations similarly and hence can continue to grow further and merge with each other. Notably, self-fusion is a unique characteristic of holothurian ossicles (Figure 3), which also extends to other echinoderms. In contrast, sponges, which also build discrete, distributed skeletons of silica or calcium carbonate, rarely display fusion between distinct branches of individually isolated spicules. Large fused skeletons, found in hexactinellids are formed uniquely using secondary silicification on existing spicules [69]. The observed cellular self-fusion in holothurian ossicle growth, thus plays a crucial role in generating topologically complex structures, but the molecular mechanisms regulating this self-fusion at the tips of the sclerocytes remain poorly understood. Thus, ossicle growth represents a distinct class of branching morphogenesis that invites further studies.

Although, previous models for biomineralization have focused heavily on bottom-up molecular components that interact with the crystalline lattices, our work highlights the complimentary role of cellular physiology in and top-down control of biomineralization. To the best of our knowledge, this is the first study of how cellular physiology directly controls and shapes topologically complex biomineral construction at these length scales. By mapping cellular scale processes such as physical wrapping and compartmentalization of the ossicle in sclerocyte cells; microtubule mediated vesicle transport; and filopodia interactions at ossicle tips, we establish a key physical framework for linking cell physiology to ossicle growth. We identify a key feedback loop, in which ossicle morphologies are patterned by sclerocytes, while sclerocytes synctium itself is shaped by the ossicle. Complete wrapping of an ossicle by a cell creates a ‘privileged space’, which is a hallmark of biomineralization [2]. The presence of a continuous, membrane-bound cytoplasmic wrapping around the ossicle forces the cell bodies to extend cytoskeletal as well as organellar components throughout the ossicle surface, which has crucial consequences for the physiology of the growth process (Figure S7G).

Despite the fact that holothurian ossicles maintain long range crystalline order, acting effectively as a single crystal (Figure S4), they are still able to execute arbitrarily curved growth of branches. How incorporation of organic matrix molecules tune local mineral curvature at molecular scale can be a subject of future studies. Similarly, if tip bifurcations are enriched in specific organic matrix molecules can be studied via molecular examination of freshly collected ossicles themselves. Furthermore, if fusion junctions leave a molecular scar can also be studied via focused ion beam milling [cite] and Nano-sims [cite] based surface imaging techniques. Several of these bottom-up processes can be subject of a future study.

We further highlight a unique similarity between processes presented here that shape ossicle growth and morphogenesis of a neuron as well as angiogenesis. The role of filopodial extensions has been previously demonstrated to be crucial in substrate-linked neuronal pathfinding as well as angiogenesis [70, 71]. In both cases (neuronal growth cones and endothelial cell tips), cells are known to reorganize cytoskeletal components in response to external gradients to execute a directed search of niches and undergo bifurcations [72, 73]. Thin filopodia in primary mesenchymal cells (PMCs) participating in skeletonization were among the first observations of sea urchin embryo development [33, 74]. These observations were further supported with experiments in isolated PMC cells in vitro [75]. The role of vascular endothelial growth factor-receptor pair (VEGF-VEGFR) in guiding PMCs toward specified ectodermal niches [76]; its conservation between sea urchins and humans [31]; the role of VEGF-VEGFR in endothelial vascularization [77]; and in vitro experiments on isolated cells [78] have inspired analogies between mammalian vascularization and PMC-regulated biomineralization of spicules. However, VEGF regulation has also been shown to orchestrate growth cone guidance and axon branching [79] and hence has been used to propose its neural origins [80]. Importantly, sclerocytes deviate structurally from vascular endothelial cells; their growth does not happen due to the proliferation of new cells, rather the same cells extend projections and undergo bifurcation at the tips, similar to those in the growth cone of neurons, while being a part of a syncytium. We believe that multi-filopodial search and tip branching is a conserved feature that has been co-opted across multiple cell types during the evolution of complexity in metazoans.

In our current model, we do not yet know what orchestrates ossicle tip bifurcation in holothorians. It is clear that the membranous projections and minimal cytoskeleton pathways play a role in facilitating tip merging, bifurcation, and budding (Figure 5). However, we do not know what explictly causes tip splitting of the mineral lattice during growth. We can rule out several physical mechanisms of branch splitting (as seen in melt solidification) as the tip splitting growth in these passive processes happens at characteristic length scales and we observe ossicles to demonstrate tip splitting at highly irregular branch lengths between successive bifurcations. The observed budding that occurs regularly at fusion sites also hints towards a process that is under cellular control rather than a passive physical mechanism. Additionally, since the timing of tip splitting sets a characteristics length scale in our problem - it also dictates the overall size of the ossicle. Addressing these knowledge gaps would require further experiments involving novel perturbations to the growth processes in living organism.

Our reduced order model captures the key idea of transport flux shaped by ossicle topology, which in turn governs growth dynamics (analogous to how a city grows, although at a different length scale) [81]. Although this is a discrete model, we do not account for crystal orientation axis/preferences due to the current lack of identified molecular players that could modulate local crystal growth. In the future, combining bottom-up molecular scale assembly with top-down approaches presented here will further strengthen multi-scale modeling of biomineralization processes.

## Conclusion

Holothurian ossicles are morphologically diverse, topologically complex calcite structures which develop via self-closing branching morphogenesis. Their development occurs within cellular, syncytial wrappings and their growth is therefore carried out through surface-restricted transport processes under tight regulation of a cellular cluster. Surprsingly, we find that soft cellular synctium assume the shape of the hard ossicle and since growth only occurs in this protected space - ossicle shape is controlled by the cellular niche. Throughout ossicle growth, accretive accumulation of biomineral as well as simultaneous expansion of the active cellular components must occur. While the mineral phase can remain stable passively, the cellular phase directly coupled to it requires continuous investment of resources by the cellular machinery for its maintenance, which limits the overall ossicle growth. This interplay could lead to the observed diversification of generated shapes through evolution of cellular processes that guide the growth of specific morphological features within an ossicle. We believe our work will inspire a new set of top-down approaches for understanding biomineralization coupled to cellular physiology, and lay groundwork for the development of in vitro systems for cell-driven self-assembly of diverse synthetic mineral geometries.

## Methods

### Spawning and larval culture

Adult *Apostichopus parvimensis* were collected off the coast of Monterey Bay, California, US, and kept in circulating seawater tanks. They were spawned during the spawning season (April to July) by injecting 5 ml of 2 nM NGLWYamide spawning peptide [82] (synthesized by Biomatik) at the anterior and posterior ends of the gonad. Sperm and mature oocytes were released by male and female animals about 45 and 90 min after the injection, respectively. Following in vitro fertilization, embryos were cultured at 14*^o^C* in UV sterilized filtered seawater, at a density of about 1 embryo per ml and within 3 l glass jars oxygenated by a motorized paddle. Filtered seawater in the glass jar was renewed every 2 or 3 days. Following water renewal, the larvae were fed ad libidum with freshly grown *Rhodomonas lens* microalgae. Auriculariae spontaneously started to undergo metamorphosis at about 3-4 weeks post-fertilization and pentactulae started to settle at the bottom of the glass jars at about 30 days post-fertilization. Settled juveniles were collected from the glass jars by being first relaxed in a 1 : 1 mix of 7.5% MgCl_2_ and seawater and then being gently detached using a paintbrush. Small batches of animals were transferred to glass bottom 6-well plates or dishes for monitoring growth and imaging. To label complete skeletons with calcein (Figure 1B), a portion of animals were grown in calcein (MP Biomedicals) labeled seawater at a final concentration of 25*µg/ml*, post the doliolaria stage, allowing calcein to get incorporated into developing ossicles [83].

### Micro-CT sample preparation and imaging

Samples were mounted using a modification of protocol suggested in [84]. Fixed samples in 4% formaldehyde in seawater were first transferred to *≥*99.8% ethanol through a serial dilution (series of seawater to ethanol ratios of 1:0, 3:1, 1:1, 1:3, 0:1). A sample-holder cylindrical tube was created by melting one end of a 20 *µ*l Eppendorf (022351656 GELoader) electrophoresis pipet tip using a hot plate until the inner diameter of the tip became almost equivalent to the width of the animal. This also resulted in melting and sealing the heated end of the tip. One or several animals were dropped from the other end with some ethanol in which they were stored. The pipette tip was struck a few times to let the animals settle to the bottom sealed end in a stable configuration. The open end of the tip was cut short to result in a 1-2 cm length cylindrical tube with one open end and animals inside. The open end was finally sealed with some capillary tube sealing clay. The sealed tube was then mounted vertically, with the heat-sealed end towards to bottom, on the flat head of a thin metal paper pin using a small drop of super glue (Loctite gel control no-drip super glue). The metal pin was then mounted on the sample holder for Zeiss Xradia 520 Versa X-ray CT and imaged using the instrument protocols. The following sets of parameters were used: Voltage- 40 kV, Power - 3W, Objective lens unit - 4x, Pixel size - 0.82*µm* for main dataset for figure 2, *<* 1*µm* for the rest, Exposure time - 15 s, Filter - air, Bin - 2, Reconstruction - AutoCB, Angle - 180 + fan, Projections - *∼* 1600. The reconstructed data was further processed, segmented and exported using ORS Dragonfly Pro (version 2022.2.0.1399). For detailed methods on data processing see supplementary information.

### Fluorescence labeling and immunohistochemistry

For membrane staining, live specimens were incubated for 15 *−* 30 min at room temperature in filtered seawater containing 1 : 1000 - 1 : 500 CellBrite Red (Biotium) lipophilic dye prior to fixation or Cell Mask Deep Red(Invitrogen) prior to live imaging. For endoplasmic reticulum (ER) staining (Figure S7J), samples were incubated with ER-Tracker Red(Invitrogen) at a working concentration of 1*µM* for 15 *−* 30 min. After incubation in live stains the samples were washed thrice with sea water before imaging. For immunohistochemistry and phalloidin stainings, we adopted a previously described protocol [85]. Animals were first fixed in 4% formaldehyde in natural filtered seawater for 30 min at room temperature. Fixed samples were then transferred to phosphate buffered saline(PBS) using serial dilution (series of sea water to PBS ratios of 1 : 0, 3 : 1, 1 : 1, 1 : 3, 0 : 1 for 5*min* each) followed by four washes in PBS containing 0.05% Tween-20 (PBST). Samples were preblocked using 10*X* Blocker BSA (Thermo Fisher Scientific, 37520) diluted to 1*X* in PBST for 1*hr*. The samples were then incubated, overnight at 4*C*, with primary antibodies diluted in PBST using anti-acetylated tubulin antibody produced in mouse (Sigma-Aldrich) and a rhodamin-coupled phalloidin (Abcam) for staining neurons and muscles (dense actin) respectively for the whole animal (Figure 1B, S8) at 0.005 U/µl. Intracellular cytoskeletal labelings were done using Anti-*α*-Tubulin*−*FITC antibody (Sigma Aldrich) and Alexa Fluor 555 Phalloidin (Invitrogen) for microtubules and actin respectively at 1:100 dilutions. Post overnight incubation, samples were washed 4 times in PBST and mounted in Vectashield Antifade Mounting Medium (H-1000-10) before on glass slide/thin glass bottom dish (Pelco, Ted Pella Inc.) before performing confocal scans. For short time scale calcein labeling, live juveniles were incubated in 25*µg/ml* concentration working solution of calcein (MP Biomedicals, 02190167-CF) in seawater. The incubation duration ranged from several hours (for Figure 5A) to overnight (for Figure 3L) and was followed by a 30*min* wash step in seawater.

### ECM live and fixed staining

Live staining of collagen was performed using collagen binding reagent Col-F(mybiosource, MBS258089) with a modified protocol from [86]. The 0.5*mg* Col-F vial contents were dissolved by adding 100*µL* DMSO to create a stock solution at 6.8*mM* and stored frozen at *−*20*^o^C*. Whole juveniles were stained at 1 : 500 in natural sea water for 15*min −* 1*hr*. Juveniles were directly imaged using confocal scanning microscopy with 488 nm excitation. A non-specific ECM labelling was achieved with ethanol dehydrated fixed samples (prepared similarly as in micro-CT sample preparation and imaging section), that were rehydrated after *>* 1 month of storage in frozen conditions (*−*20*C*) using serial dilution into 1X PBS (series of ethanol to PBS ratios of 1:0, 3:1, 1:1, 1:3, 0:1 for 5 mins each). Subsequently the samples were stained with Hoechst 33342 1mg/ml stock solution at 1:500 dilution. Although this protocol was sed for labelling nuclei in samples, this resulted in ECM labelling throughout the animal tissues whose specificity could not be determined. The stained fibers overlapped the dense fibers visible in ECM substrate near ossicle hinting towards their identity as collagen bundles observed in TEM images (Figure S6C,D). Similar protocol was recently reported for staining collagenous tissues in anhydrous samples using Fast Green [87].

### Live brightfield imaging

Multiple imaging configurations were used for live ossicle time lapse imaging. Images shown in Figure 3J were obtained by embedding the animal in a droplet of 0.1 *−* 0.5% agarose in a glass bottom dish filled with seawater. This slightly constrained the animals movement, while allowing the diffusion of seawater allowing unconstrained ossicle growth. The animal was then imaged at 40X magnification, 3 fps using a polarization filter for 5 days. This approach required manual processing to track specific ossicles on the body at different time points time. Images shown in Figure 3F and Figure 3G were obtained by gently compressing the animal with a glass coverslip with PDMS/imaging spacer in between to reduce the amount of compression appropriately. This allowed continuous imaging of single ossicles for shorter intervals of time (on the order of several hours rather than days). Animals were relaxed or temporarily immobilized in a 1 : 1 mix of 7.5% MgCl_2_ and seawater for imaging of cellular processes at higher temporal resolution, including datasets in Figures 5G,H,K. For Figure 5K, a live juvenile was partially dissociated using collagenase type IA (at 0.5 mg/ml dissolved in calcium and magnesium free sea water made using cold spring harbor protocols) and ossicles within dissociating tissue were imaged directly, or after isolating them through gentle pipetting. All brightfield live imaging was performed using Nikon Ti2 Eclipse inverted microscope (with NIS Elements software, Nikon objective lenses: 4x/0.13 air Plan Fluor, 10x/0.30 air Plan Fluor, 20x/0.75 air Plan Apo, 40x/0.13 oil Plan Fluor, 60x/1.49 oil Apo TIRF, Camera: Photometrics Prime 95B 25mm) or customized tracking microscopes built using the SQUID platform [88].

### Confocal imaging

Imaging of fluorescently labelled animals/tissues was performed using a Zeiss LSM700, LSM780 or LSM 880 confocal microscopes and multiple versions of Zen Black softwares coupled with the microscope setups were used for acquisition. The following Zeiss objective lenses were used: 10x/0.3 air EC Plan-Neofluar, 20x/0.8 aair Plan-Apochromat, 40x/1.3 oil EC Plan-Neofluar and 63x/1,4 oil Plan-Apochromat. Multichannel acquisitions were obtained by sequential imaging and excitation - emission spectra and hardware configuration were set using smart setup (quality, best signal) option within Zen black software. Datasets were acquired using Confocal optical sections were compiled into maximum intensity z-projections using ImageJ – Fiji v.1.54f [89].

### Transmission electron microscopy sample preparation and imaging

We utilized a modified version of protocol from Li et al [90] for making resin embeddings for samples. All steps were performed at room temperature (RT, *∼* 25*^o^C*) unless explicitly stated. Fixed animals were washed serially PBS as mentioned before and were stained with Safranin-O, stabilized in 1% low melting point agarose, and then cut into 1mm cubes to remove excess agarose gel. Samples were rinsed (thrice for 5*min* each) in 0.1*M* sodium cacodylate buffer (pH 7.4) and immersed in 20mM glycine solution in the same sodium cacodylate buffer for 20 minutes at 4*^o^C*. The samples were then rinsed again three times with sodium cacodylate buffer. The samples were fixed in secondary fixatives 5*mL* of 4%*wt/vol* osmium tetroxide aqueous solution mix with 500*µL* of 20%*wt/vol* potassium ferricyanide and 4.5*mL* of 0.1*M* sodium cacodylate buffer for 1*hr* at 4*C*. Samples were rinsed twice in sodium cacodylate buffer and left at RT to warm up for 20 minutes and then briefly rinsed in molecular biology reagent grade water (MBR water). Samples were then stained in 1% thiocarbohydrazide in MBR water for 20 minutes. Samples were rinsed thrice for 5*min* each in MBR water, followed by 2% osmium tetroxide aqueous solution for 1*hr* in dark. Samples were again rinsed twice in MBR water, and stained in 4% uranyl acetate overnight. Samples were rinsed thrice in MBR water, and stained in 0.15% tannic acid in MBR water for 3*min*. After rinsing twice in MBR water, samples were transferred to ethanol through a serial dilution (10min each in 30%, 50%, 70%, 90%, 95%, 95% ethanol in MBR water) followed by pure ethanol washes (thrice, 10 min each). Samples were then placed into 100% propylene oxide (post one wash in the same for 5 min). This was followed by infiltrating sample with Embed 812 resin (EMS, Hatfield, PA, USA) through serial dilution (ascending gradient of resin: propylene oxide mixture, 1 : 3 for 2*hrs*, 1 : 2 for 2*hrs*, 1 : 1 overnight, 2 : 1 for 3*hrs*, whole resin for 4*hrs*), while rocking. The EMbed 812 resin was made by mixing 100*mL* of EMbed 812 with 45*mL* of DDSA, 60*mL* of NMA and 6.15*mL* of BDMA, to make a harder block more suitable for TEM sectioning. The resulting resin blocks were chipped off until the embedded sample was exposed and then ultrathin sections (80*−* 100*nm* thick) were made and post stained with uranyl acetate and lead citrate. The sections were then imaged on Transmission Electron Microscope JEOL JEM1400 using appropriate settings at various magnifications.

### SBF-SEM sample mounting and imaging

Samples were embedded in epoxy with the same method as used for TEM and shipped to University of Illinois Urbana-Champaign for SBF-SEM. A protocol similar to the one previously utilized by the facility was used [91]. Before initiating block face imaging, epoxy-embedded samples were mounted to aluminum pins (Gatan Inc., Pleasanton, CA, USA) using silver epoxy (Ted Pella) and sputter coated with a thin layer of Au/Pd. Serial block-face imaging was performed using a Sigma VP (Zeiss) equipped with a Gatan 3 View system (model: 3View2XP) and a nitrogen gas injection manifold (Zeiss model 346 061-9002-200). Thick slices were first chipped off to reach the desired imaging site with a lateral view of ossicle cross section which was then imaged at 2.0*keV*, using 50*nm* cutting intervals, 10.0*nm* pixel size (12*k ×* 12*k* pixels), beam dwell time of 1.0*s* and a high vacuum chamber pressure of *∼* 5 *×* 10^.3^*mbar*. The SEM images were then stacked, processed, segmented and exported as 3D views/videos using ORS Dragonfly Pro (version 2022.2.0.1399).

## Acknowledgements

We thank all members of the Prakash Lab for open-ended discussions that significantly improved the quality of the work. In particular, the authors acknowledge Ray Chang for insightful discussions and help with the TEM sample preparation and Rebecca Konte for discussions on figure graphics. We also thank Neeraja Abhyankar for her feedback on modeling and figure preparation.We thank the following central facilities at Stanford University – Cell Sciences Imaging Facility Shriram Center for Bioengineering and Chemical Engineering and Beckman Center for Molecular and Genetic Medicine (Youngbin Lim, Kitty Lee, Cedric Espenel, David Lenzi), TEM facility (John Perrino, Ibanri Phanwar-Wood and Jon Mulholland), Micro-CT facility (Arturas Vailionis) and SBF-SEM facility (Kingsley Boateng) at University of Illinois at Urbana Champaign. We thank Prakash lab admin Jasmine Desiderio for timely availability of experimental resources. We are thankful to the Monterey Abalone Company for annually sourcing adult *A. Parvimensis* for the spawns. The authors also gratefully acknowledge the following funding sources: Pranav Vyas (Stanford Graduate Fellowship, BioX - Stanford Interdisciplinary Graduate Fellowship, Siebel Fellowship), Charlotte Brannon (Training grant from NIH Cellular and Molecular Training Grant, NIGMS grant number 5T32GM007276 and National Science Foundation Graduate Research Fellowship Program), Manu Prakash (CZI BioHub, HHMI Faculty Fellowship, Schmidt Foundation, Moore foundation, NSF CCC Grant). The authors declare no conflict of interest.

## Author credits

PV, CB and MP designed the research. LF and CL provided critical resources and cultures. PV, CB, LF and MP collected the data. PV, CB analyzed the data in discussion with MP. PV developed the analytical model in discussion with MP and CB. PV, CB and MP wrote the manuscript. All authors commented and edited the final version of the manuscript.

