## Supplementary text file for "Algorithmic construction of topologically complex biomineral lattices via cellular syncytia"

### Supplementary Information: Algorithmic construction of topologically complex mineral lattices via cellular syncytia

February 20, 2024

**Keywords:** Holothuroidea, Biomineralization, Morphogenesis, Self-closing branching networks, Syncytium

#### Supplementary Figures S1-S9

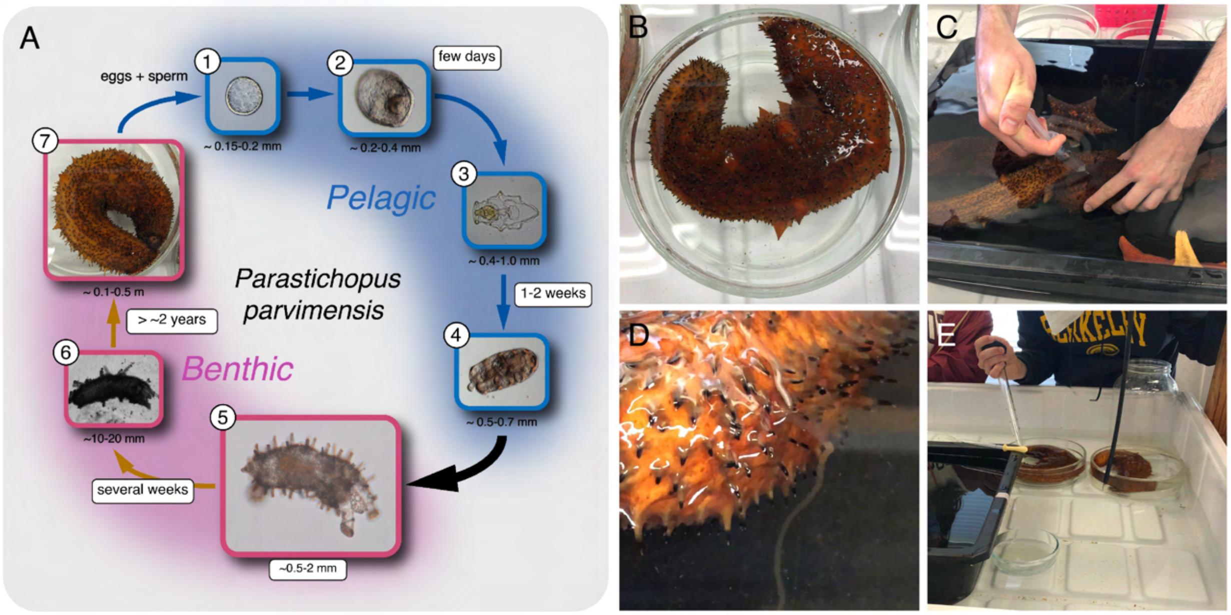

**Figure S1: Life cycle of *A. parvimensis* and spawning methods** (A) *A. parvimensis* or warty sea cucumbers are epibenthic detritivore echinoderms, found along the coast of California. Separate separate male and female individuals aggregate and perform broadcast spawning during March to July every year, releasing eggs and sperms in the ocean and that get fertilized externally (1). Once fertilized, the embryo undergoes a series of cell divisions, followed by gastrulation within the next few days (2). The gastrulated swimming larvae develop further into swimming Auricularia which filter feed swimming planktons from the ocean (3). Within a week or two the Auricularia metamorphoses into Doliolaria stage. The Doliolaria develops into the pentacula stage within a few days (4 depicts this transition) which is accompanied by simultaneous seeding and growth of ossicles all over the animal body and transition into a benthic detritivorous life style. The ossicles develop and the pentacula transforms into the earliest juvenile within a few weeks (5-6) which grow over several years to reach sexual maturity. New ossicles are added throughout the growth of the animal. (B-E) Spawning peptide induced aritificial spawning of adult *A. parvimensis*. (C) Peptide is injected into the body cavity of sexually matured adults collected during the spawning season. (D) The injection induces sperm and egg shedding within 1-1.5 hrs which are then collected using pipettes and stored separately. The gametes are later combinedin glass beakers to induce fertilization.

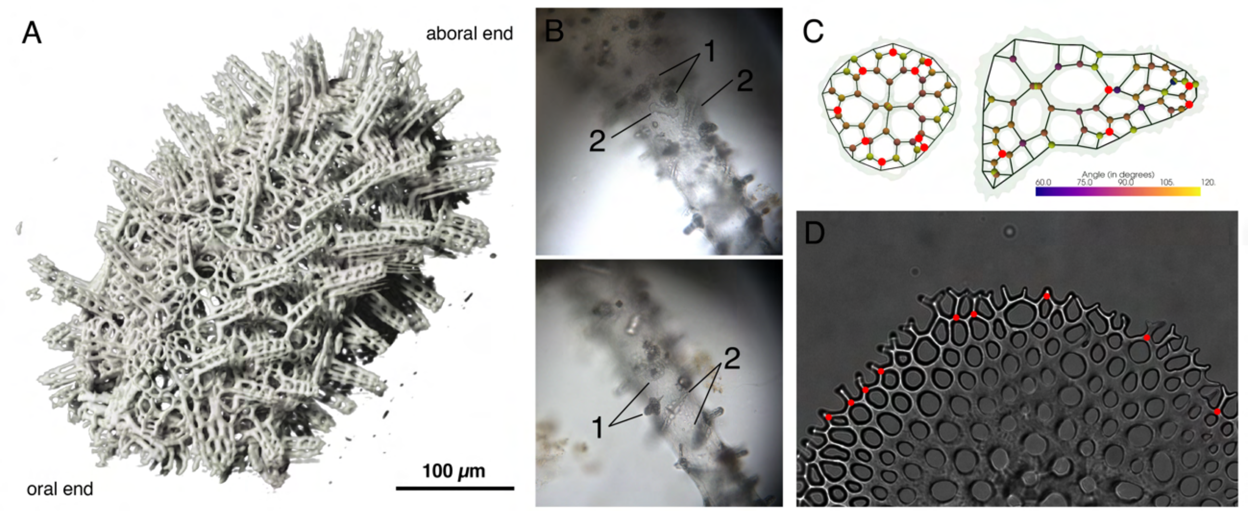

Figure S2: (A) An oblique view rendering of micro-CT scan of juvenile *A. parvimensis*. (B) Observation of simultaneously growing distinct ossicle types, 1 - pillared table, 2 - elongated plate, in identical tissue neighborhood in a tube feet. This suggests that ossicle types are not necessarily prescribed by the local tissue neighborhood, and agency of sclerocytes building them must be important. (C) Identification of putative budding sites in already developed ossicles as nodes with two incoming edges and one outgoing edge; directions identified with increasing node depth values (see supplementary methods) (D) Several putative budding nodes identified in a foot pad ossicle, where the node is almost equidistant from the nodes that initiate branches which fuse to create the concerned node.

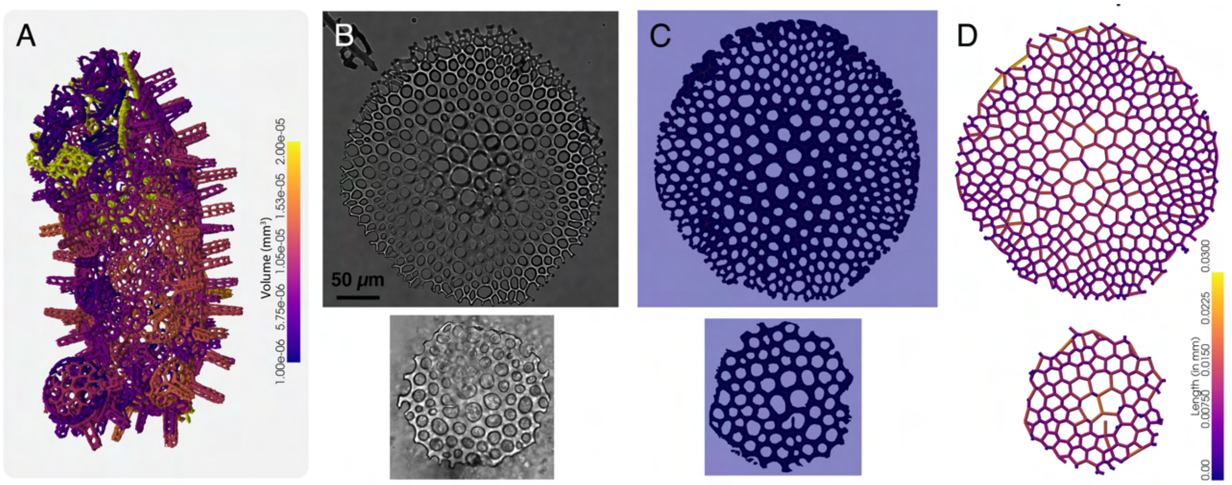

Figure S3: (A) Map of ossicles color labeled based on their volume. (B) Adult foot pad ossicles used for geometric and topological computations in Figure 2. (C) Threshold based segmentation and skeletonization. (D) Map of all edges of ossicles color labeled based on their lengths.

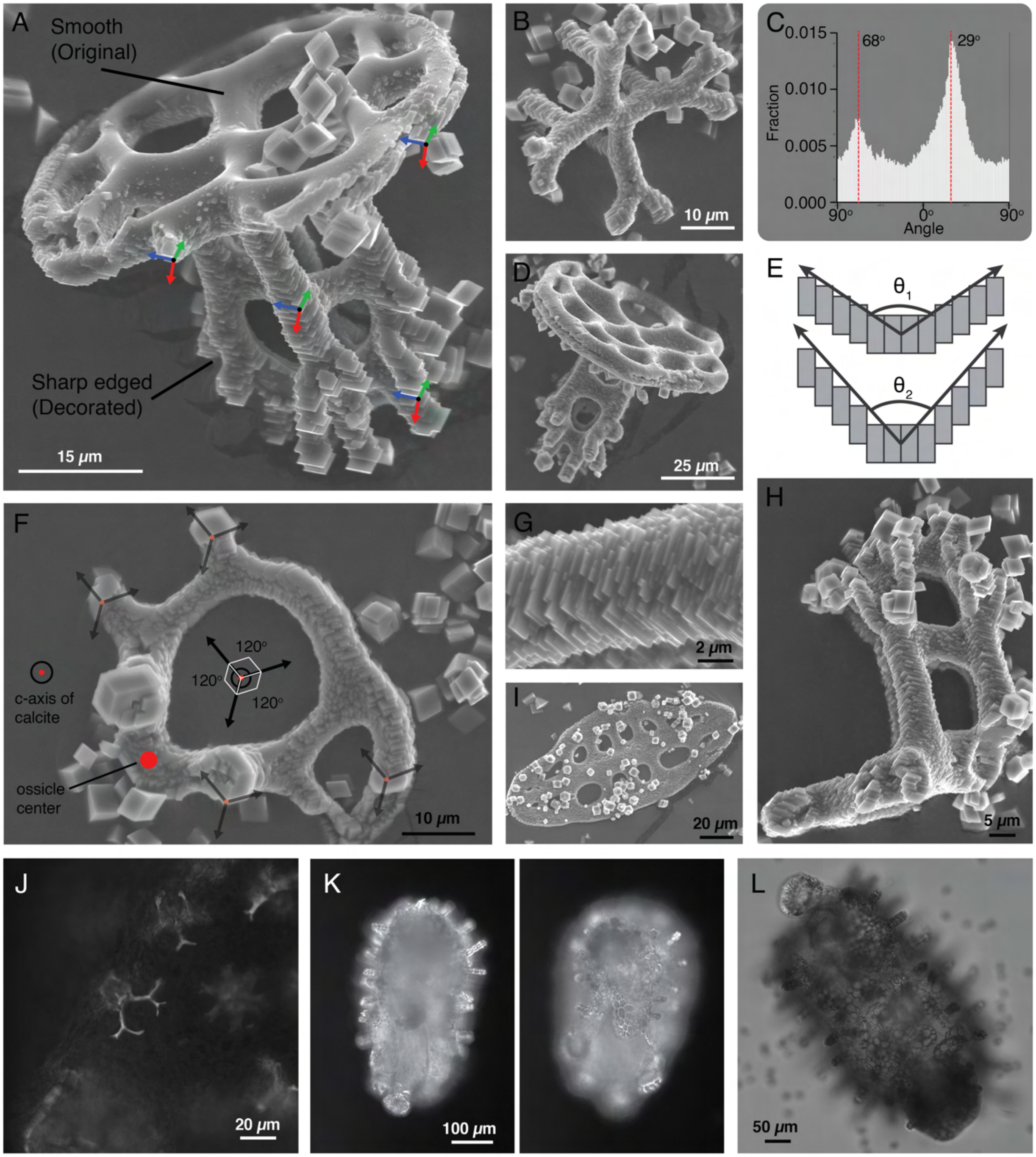

**Figure S4: Long range preservation of crystalline order in ossicles** (A, B, D, F, G, H, I) SEM micrographs of isolated *A. parvimensis* ossicles with epitaxially grown calcite crystallites (sharp rectangular edges) with smooth ossicle surface as a template (see supplementary methods). (C) Direct visual interpretation and quantification reveals preservation of crystalline order across the entire ossicle geometry. The facets of precipitated portions remain parallel throughout the ossicle, irrespective of curved edges or vertical extensions in the ossicle shape (arrow triplets in panel A). The image in panel B representing an early stage ossicle, was thresholded and segmented to identify bright edges representing the facets. We plotted binned angular orientation of these edges and observed two distinct peaks representing the most visible facet orientations.

(E) Within ossicles, this could be achieved through accretive deposition of relatively displaced precursor material boluses (or crystallites [1]), which could be stabilized by the organic matrix molecules, resulting in curved geometries. (F) A broken quarter portion of a pillared table ossicle oriented in a top view. Most of the grown crystallites display bright edges, representing facet orientations, in directions separated by $\sim 120^o$. In rhombohedral calcite symmetry, such an axis resembles the c-axis of calcite, which in this case aligns with the vertical growth direction of the pillared table (perpendicularly away from the body surface). This observation was consistent across $\sim 5$ distinct pillared table ossicles. Such a conserved orientation of crystal during ossicle growth has been previously proposed in other holothurian species [1] and very recently for sea star ossicles [2]. However, how this remarkable feat is achieved by cells across the animal body is in interesting subject for future studies. (G) Zoomed in view of one of the branches. (H) Side view with exposed surfaces from broken branches decorated as well, showing that the crystalline order penetrates beneath the ossicle surface as well. (I) Top view of an elongated plate showing precipitated growth with c-axis pointing vertically outward from the page, similar to the case of pillared table ossicles. In all the images for precipitation induced growth, there are many large crystals scattered across images. These crystals grew independently in the solution and then settled on the ossicles, rather than growing on the ossicle surface as templates. They represent noise in the system and need to be ignored while evaluating orientation of crystal facets. (J) The preservation of crystalline order across the entire structures allows the ossicle to interact with light like a single crystal and display birefringence under polarized light. (K) Whole animal view displaying birefringence across all ossicles under polarized light. (L) DIC microscopy view of an entire juvenile, for a comparison to the polarization one. Here individual ossicles do not shine up due to birefringence.

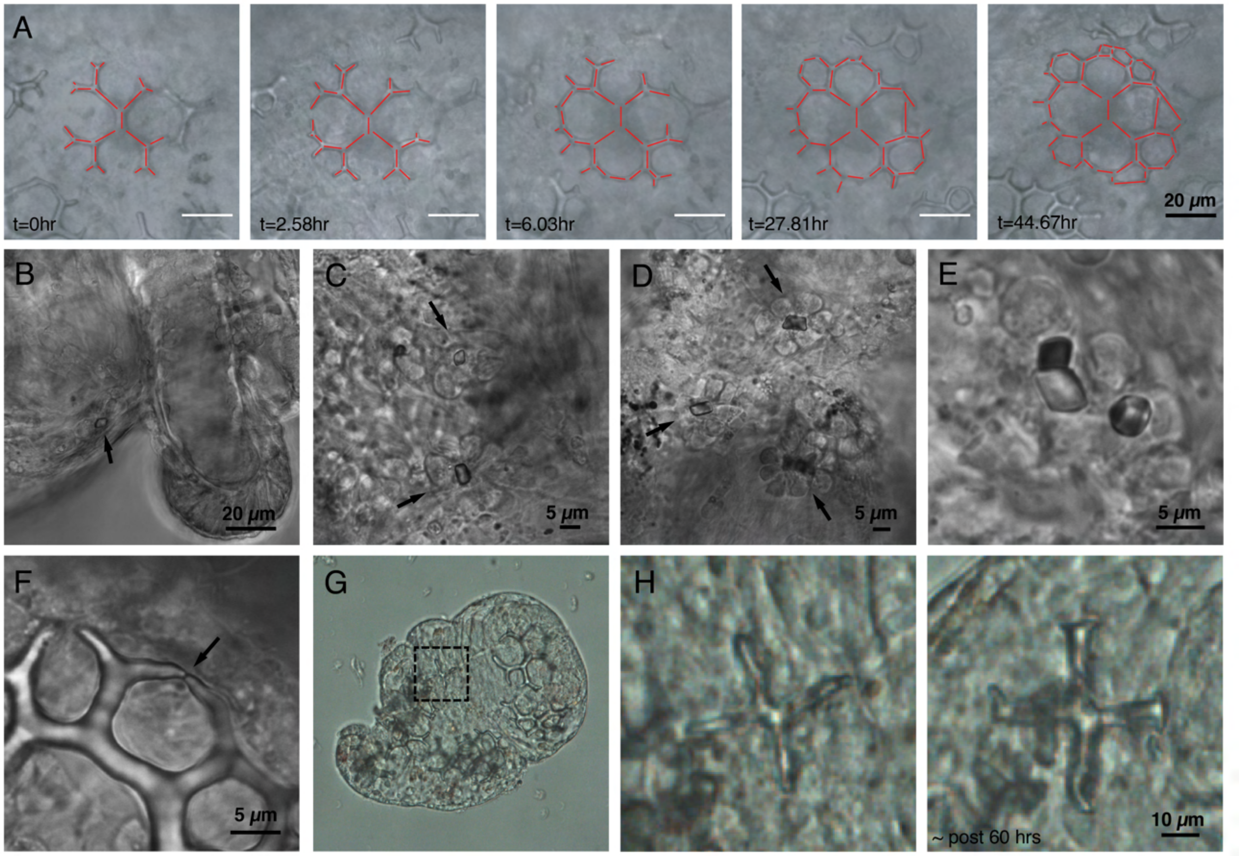

Figure S5: (A) A second growth series depicting asymmetric and asynchronous growth in pillared table ossicles, similar to that in Figure 3M. (B-E) Seed crystals and associated sclreocyte clusters in different contexts within the body. Panel E shows differently oriented rhombohedral seeds (arrows) within the sclerocyte syncytium. (F) A snapshot of an instance where two biomineral tips are about to merge (arrow). (G) An isolated tube feed from a juvenile *A. parvimensis*, slightly squished under a coverslip and being imaged continuously. (H) Even within the isolated foot, the an ossicle displayed slow growth over a period of 60 hours, suggesting that sequestration of mineral precursors from sea water happens locally within the tissues and is not packaged centrally and then circulated within the entire animal body.

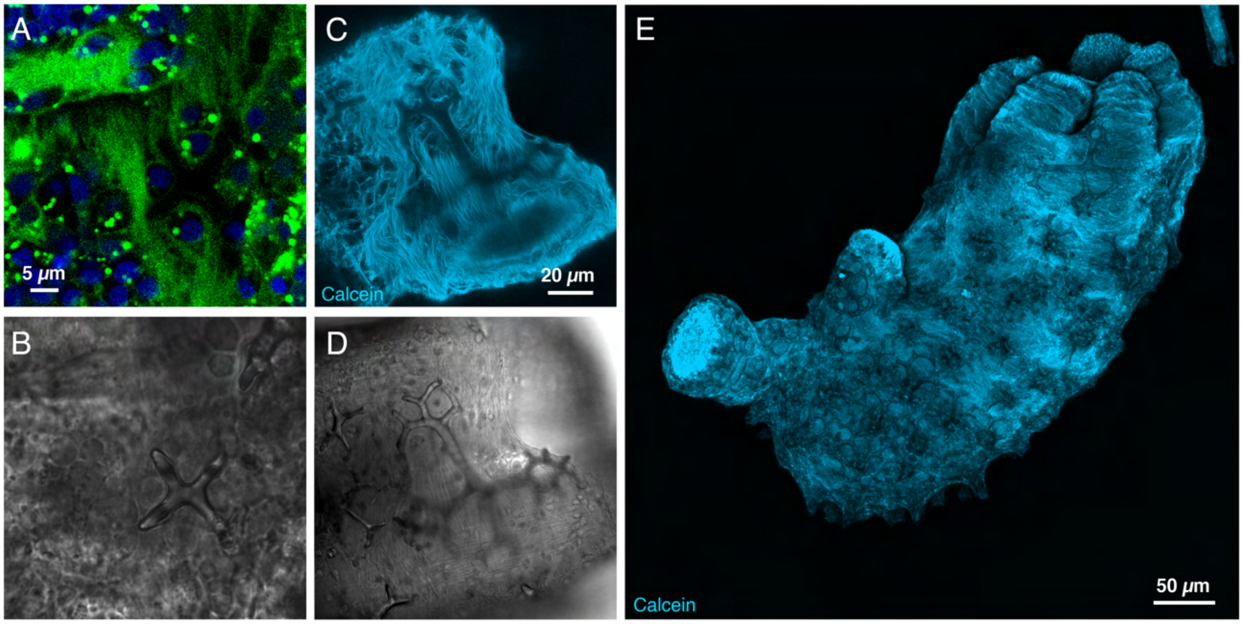

**Figure S6: Direct imaging of ECM surrounding growing ossicles.** (A) Live Col-F stain (Green) to stain collagen and Hoechst stain (Blue) for nuclei were used to reveal arrangement of collagen fibrils near growing ossicles. This image is an additional data set complementing Figure 4H. Panel B shows the transmitted photo multiplier tube (T-PMT) channel image for the same. (C) Confocal scan showing non-specific labeling of ECM near a growing ossicle (see methods). Panel D shows the transmitted photo multiplier tube (T-PMT) channel image for the same. (E) Maximum intensity projection of confocal scan of non-specific ECM labeling for a whole juvenile. All scans reveal close association of ECM with the ossicles but lack of any organization that could act as template for the growth of the ossicle.

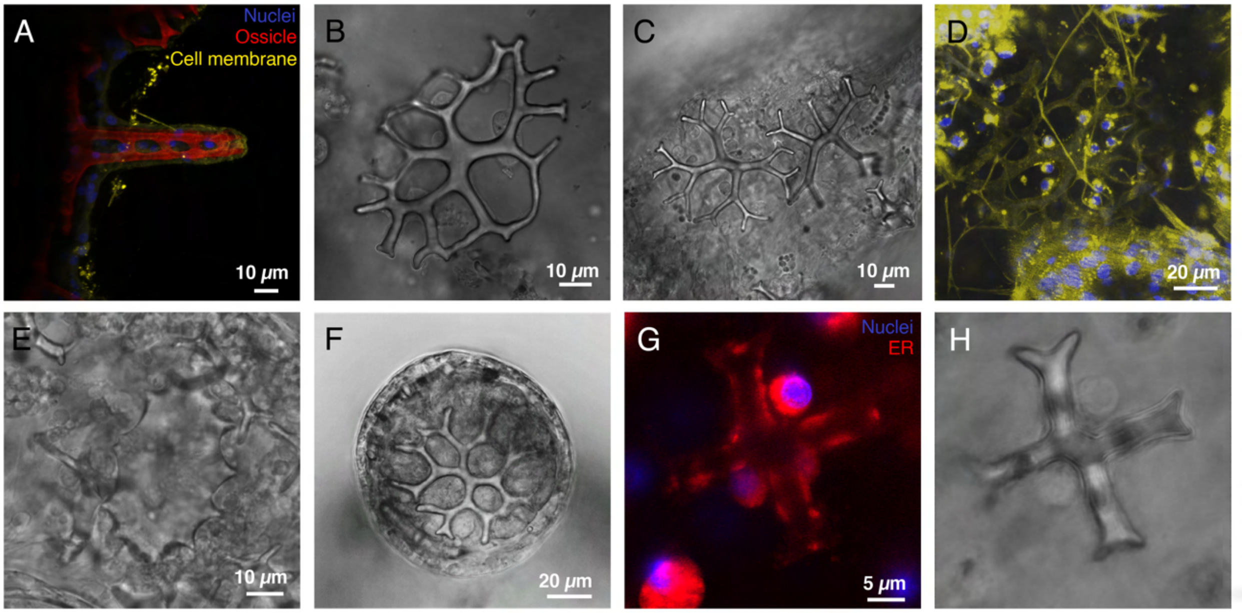

Figure S7: (A) Maximum intensity projection of confocal scan of a lateral view of vertical growth of a pillared table ossicle. Just like planar growth, the vertical growth also displays decreasing hole and branch sizes. (B) Ossicle with topologically trapped sclerocytes hanging on its surface after being separated from the body wall using collagenase treatment. (C) DIC imaging of early stages of growth of front plate ossicles (D) A confocal scan of front plate ossicles with cell membrane (yello) and nuclei (blue) labels displaying associated sclerocytes spread further apart on the structure as compared to pillared table ossicle. (E) Larval ossicle remnant remaining near the rear end of a young juvenile. (F) An early stage foot pad ossicle embedded within the foot pad tissue. (G) Early cross stage pillared table ossicle, with ER (red) and nuclear (blue) stain showing ER distributed in small patches throughout the syncytial cytoplasmic sheath wrapping the ossicle. Panel H shows the transmitted photo multiplier tube (T-PMT) channel image for the same. Distribution of many organelles, especially Endoplasmic Reticulum (ER) and Golgi Apparatus (GA) have been shown to play a role in stabilizing branched dendrite morphologies, especially in the case of neurons. [3, 4]. These properties make growth cones a great analogy to growing tips in ossicles.

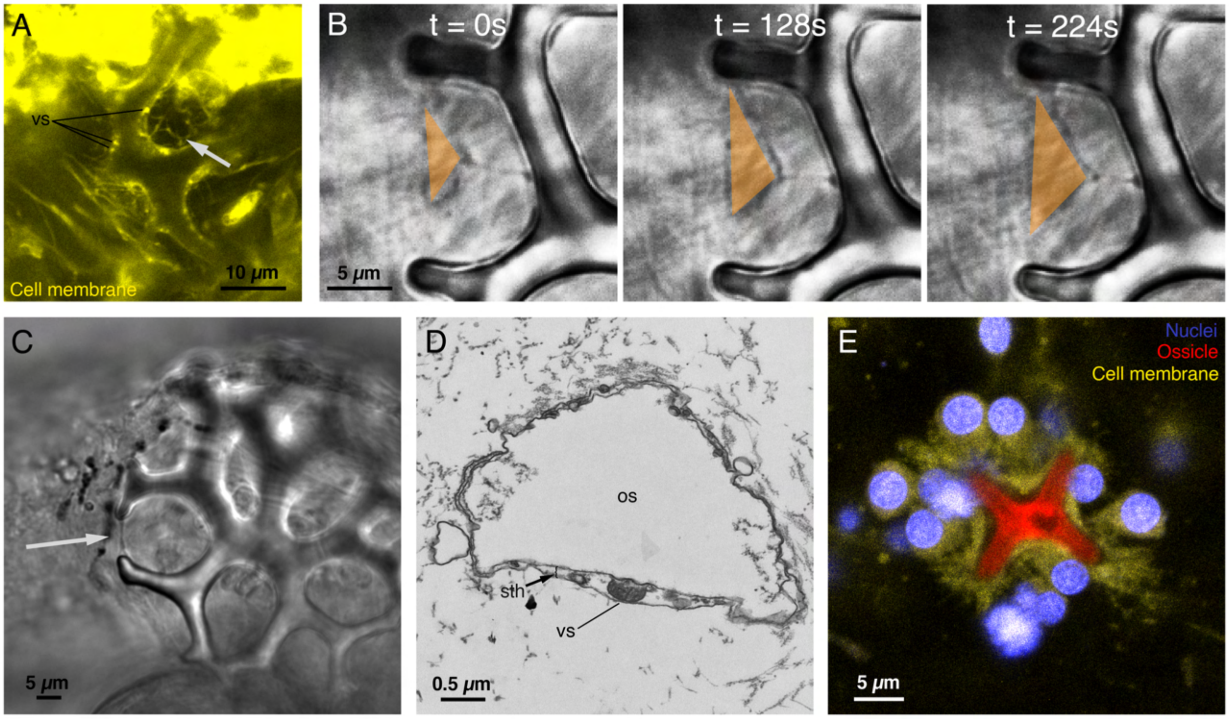

**Figure S8: Membrane and vesicle activity during ossicle growth.** (A) Live fluorescence labeling of membranes (yellow) highlighting membrane bound vesicles (vs) in the cytoplasmic sheath as well as thin membranous projections within ossicle hole region (arrow). (B) DIC imaging of a region between two extending tips shows fluctuating membrane threads in a triangular configuration (orange). Such miscellaneous membranous structures can be observed between branches throughout the ossicle, sometimes accompanied with motile boluses transported through them. The changing area of the triangular region depicts fluctuation in membranes (see supplementary video SV6) (C) DIC image of an ossicle from a live juvenile displaying a membrane thread between two approaching tips (arrow) (D) TEM image of a cross section of an ossicle branch displaying a putative merging event of a vesicle (vs) with the inner membrane. os - ossicle, sth - cytoplasmic sheath. (E) Multi-channel confocal scan of an early stage ossicle displaying thin membranous projections at the tips (arrow).

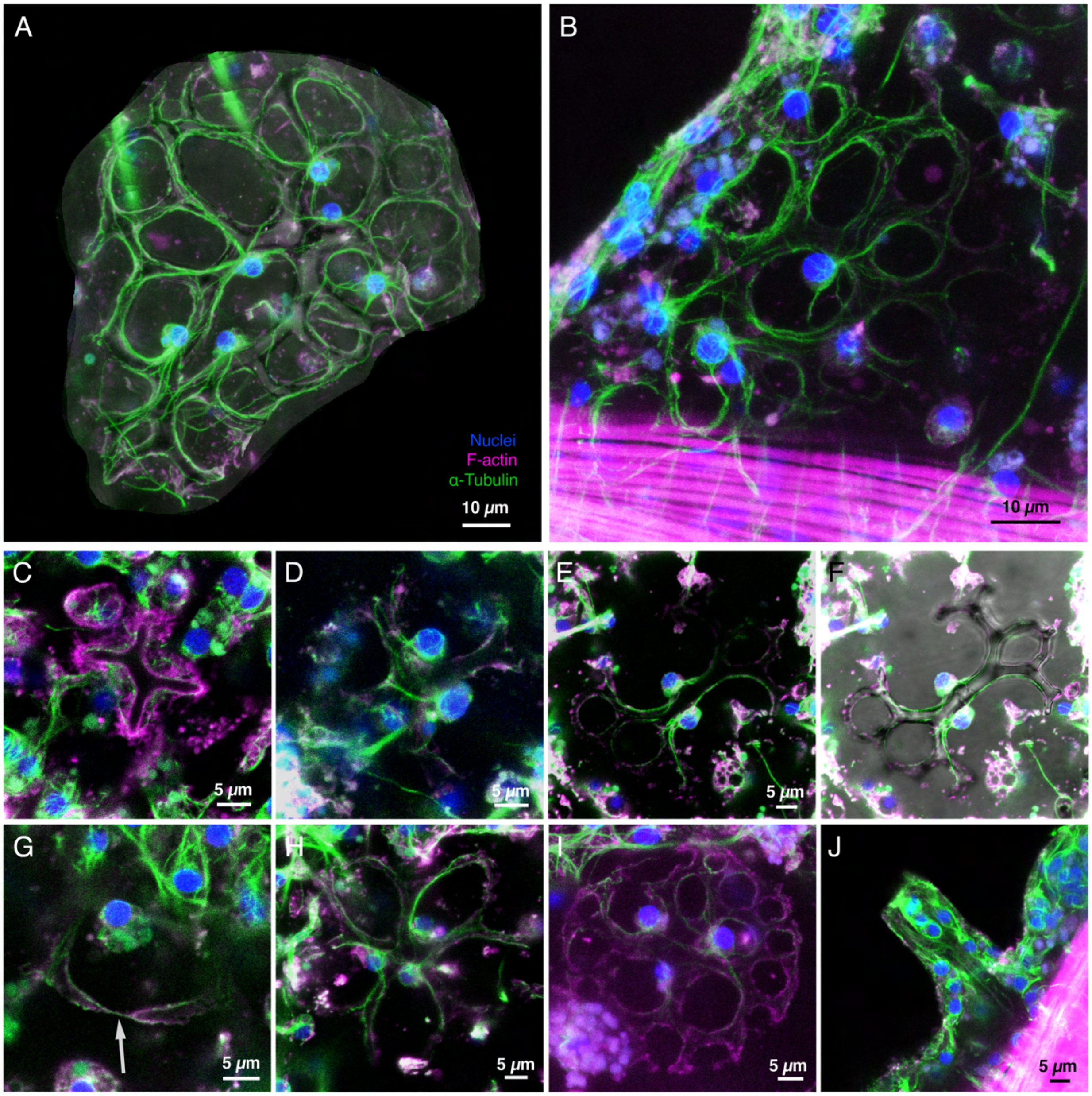

**Figure S9: Cytoskeletal organization within syncytium at different stages of ossicle growth** (A) A maximum intensity projection of confocal z-stack of a fixed front plate ossicle in which each plane is cropped to only cover the ossicle region. This allows removal of signals from other layers of the tissue and the elaboration of complete cytoskeletal network withing the ossicle-sclerocyte syncytium. The alpha tubulin and f-actin signals are overlaid on top of T-PMT channel signal from one of the planes to visualize the ossicle contour with respect to the cytoskeleton. (B) A stack projection for another front plate ossicle with ossicle mineral completely dissolved due to long (>1 day) incubation in 1x PBS. This reveals cytoskeleton from the entire syncytium, even from the back of the ossicle, which usually get hidden due to optically opaque ossicle. (C) A stack projection early stage ossicle (dark region), displaying numerous actin rich filaments at the tips. (D) A stack projection for an ossicle displaying first bifurcations of tips. (E) A stack projection for an intermediate stage front plate ossicle growing with just 2 cells, which is rarely observed. This leads to a dramatically distinct overall geometry, yet the characteristic features of branching bifurcations are still preserved. (F) Same as E but overlaid on top of T-PMT signal.

(G) A zoomed in view of membrane thread (arrow) created by two tips that are about to fuse. The thread displays both alpha-tubulin and f-actin signal. (H) A stack projection for a pillared table ossicle with pentaradial symmetry displaying some of the first fusions. (I) A stack projection for a fully formed pillared table ossicle. The weak signal is due to higher penetration depth required to image in region near the vertical pillar. But the ossicle displays both alpha tubulin and f-actin signals on all the branches of the ossicle. (J) A stack projection for a the side view of a pillared table ossicle. Cytoskeletal labeling specific to the syncytium is difficult to isolate due to overlapping signals from the epidermal tissue.

#### One dimensional growth analytics

Here we present the 1D model of directed growth - modeling microtubule supported vesicle transport of mineral components. Assume that mass is being injected at a point at a rate $\dot{m}_o$, say the origin, and the ossicle grows linearly along the positive x-axis. The decrease in mass rate due to fractional dropping can then be written as -

$$\frac{d\dot{m}}{dx} = -\eta\dot{m} \tag{1}$$

Here $\eta$ is the fraction total mass dropped when it traverses over a unit distance. An alternative extreme of this could be constructed using an assumption that a constant amount of mass, $\eta'$, gets dropped every unit distance, in which case the mass rate change equation would become -

$$\frac{d\dot{m'}}{dx'} = -\eta' \tag{2}$$

Integrating these, we get the expressions for local mass rate with distance as -

$$\dot{m} = \dot{m}_o e^{-\eta l} \tag{3}$$

or for the constant drop rate case -

$$\dot{m'} = \dot{m}_o - \eta' l' \tag{4}$$

Now let's consider the length growth of a branch due to such a mass rate distribution. In the continuum limit we can write the time rate of increase in length of the branch as -

$$\frac{dl}{dt} = \mu\dot{m}_t \tag{5}$$

Here $\dot{m}_t$ is the mass rate available at the tip of the branch and $\mu$ is a physical factor that decides the branch length increase due to a unit mass accumulation at the tip. This can be calculated for a spherically capped tip with a diameter that remains constant throughout the growth and a constant material density that also takes into account the conversion from a precursor stage to the final crystallized biomineral. Assuming $\rho$ as the density of biomineral in the final form, and $w_o$ as the constant inital tip diameter, we get $\mu = \frac{4}{\rho \pi w_o^2}$. Using this, the length of the branch can be evaluated as a function of time as follows -

$$l = \frac{1}{\eta} ln(\eta \mu \dot{m}_o t + 1) \tag{6}$$

Similarly for the constant drop rate case, we get -

$$l' = \frac{\dot{m}_o}{\eta'}(1 - e^{-\eta' \mu t'}) \tag{7}$$

For the fractional drop case, the logarithmic function grows at a slowing pace, but never terminates, unless we impose a threshold mass necessary for extension of length. However, for the constant drop rate case, the maximum achievable length becomes $l_{max} = \frac{\dot{m}_o}{\eta'}$.

Using geometrical conversion, we can write down the rate of thickness ($w$) increase as -

$$\frac{dw}{dt} = \frac{-2}{w \rho \pi} \frac{d\dot{m}}{dx} \tag{8}$$

For the case of fractional dropping, this reduces to -

$$\frac{dw}{dt} = \frac{2}{w \rho \pi} \eta \dot{m} = \frac{\mu w_o^2}{2w} \eta \dot{m} \tag{9}$$

And similarly for the case of constant drop rate, we get -

$$\frac{dw'}{dt'} = \frac{2}{w' \rho \pi} \eta' = \frac{\mu w_o^2}{2w'} \eta' \tag{10}$$

Assuming that the tip initiates with a diameter $w_o$ at $t = 0$, we can integrate these to get the temporal growth of the width at a specific position as -

$$w = w_o \sqrt{1 + \mu \eta \dot{m}_l t} \tag{11}$$

Where $\dot{m}_l = \dot{m}_o e^{-\eta l}$ is the mass rate available at a distance $l$ away from the site of injection, which can be considered to be constant for a fixed position. Similarly, we can integrate for the case of constant drop rate to get -

$$w' = w_o\sqrt{1+\mu\eta' t'} \tag{12}$$

For both the cases, the thickness grows as a function of $\sim \sqrt{t}$.

In order to evaluate the shape profile of the branch as a function of time, we can first evaluate the time of inception of a point as the branch grows as follows -

$$t(x) = \frac{1}{\eta\mu\dot{m}_o}(e^{\eta x} - 1) \tag{13}$$

$$t'(x) = -\frac{1}{\eta'\mu} ln\left(1 - \frac{\eta' x'}{\dot{m}_o}\right) \tag{14}$$

Substituting these into the functions for width with time gives functions of width profile, which is dependent on both position and time -

$$w(x,t) = w_o e^{\frac{-\eta x}{2}}\sqrt{1+\mu\eta\dot{m}_o t} \tag{15}$$

$$w'(x,t) = w_o\sqrt{1 + ln\left(1 - \frac{\eta' x'}{\dot{m}_o}\right) + \mu\eta' t'} \tag{16}$$

### Relevant scales and quantitative numbers associated with biomineralization in sea cucumbers

| | | |
|---|---|---|
| | **Spatial scales** | |
| 1. | Cell size, $s_c$ | $\sim 7\mu m$ |
| 2. | Cell volume, $V_c$ | $\sim 1400\mu m^3$ |
| 3. | Nuclear size, $s_n$ | $\sim 3\mu m$ |
| 4. | Nuclear volume, $V_n$ | $\sim 100\mu m^3$ |
| 5. | Cytoplasmic volume, $V_{cyt}$ | $\sim 1300\mu m^3$ |
| 6. | Ossicle size, $s_o$ | $\sim 50\mu m$ |
| 7. | Branch width, $w_b$ | $\sim 5\mu m$ |
| 8. | Mean cytoplasmic sheath width, $w_{cyt}$ | $\sim 0.25\mu m$ |
| 9. | Ossicle Volume, $V_o$ | $\sim 5000\mu m^3$ |
| | **Temporal scales** | |
| 1. | Transcriptional time scale (mammalian), $t_{tr}$ | $\sim 10 - 100nts/s$<br>$\sim 10min/gene[300aa]$ [5] |
| 2. | Translation time scale (mammalian), $t_{tl}$ | $\sim 10aa/s$<br>$\sim 1min/protein[300aa]$ [5] |
| 3. | Speed of vesicle transport over microtubule in free cytoplasm, $v_{ves}$ | $\sim 1\mu m/s$ |
| 4. | Observed bolus movement speeds, $v_{bol}$ | $\sim 0.1\mu m/s$ |
| 5. | Actin-induced tip membrane activity time scale, $t_{act}$ | $\sim 100s$ [6] |
| 6. | Tip growth speed, $v_{tip}$ | $\sim 1\mu m/hr$ |
| | **Other numbers** | |
| 1. | Ossicle Mass, $M_o$ | $\sim 13550pg \sim 13550 \times 10^{-12}g$ |
| 2. | Number of cells in a cluster, $N_c$ | $\sim 6$ cells |
| 3. | Calcite density, $\rho$ | $\sim 2.71g/cm^3$ |

Table 1: Relevant physical scales and quantitative numbers for biomineralization in sea cucumbers - either measured directly in our experiments or estimated from literature

### A note on protein incorporation rates

Rate of volume increase during ossicle growth, as estimated in Figure 3K is $1.486 \times 10^{-7} mm^3/hr$. Known calcite density, if assumed to be unchanged due to organic matrix incorporation is $2.71g/cm^3$. Using reported estimates [7] of organic matrix percentage within sea urchin biomineral as $< 1\%$ by mass and the complex proportions of proteins, proteoglycans and polysaccharides in the organic matrix [8], we choose an approximate weight percentage of proteins to be 0.5%. Using these we can calculate the total mass accretion rate to be $402.71pg/hr$, and the protein mass incorporation rate to be $2.01353pg/hr$. Assuming an average translation rate of $10^5$ proteins/$s$ [9], similar to that of a mammalian cell, an average protein size of 472 amino acids [10], and an average amino acid mass to be $110Da$ or $1.82659 \times 10^{-10} pg$, we can estimate protein mass production rate in a single cell to be $31.04 pg/hr$. For 6-7 cells, the total protein mass production rate can reach around $200 pg/hr$, which is around 2 orders of magnitude higher than the estimated value from the protein mass incorporation rate in the biomineral, suggesting that a very small percentage of all proteins produced in a cell are used for ossicle matrix.

#### Estimating number of ossicles on an adult *A. parvimensis*

From the micro-CT datasets, we counted the number of ossicles for a juvenile of about $\sim 500\mu m$ length and $\sim 200\mu m$ width to be about 200. For the largest possible size of *A. parvimensis*, assumed as a cylinder of $\sim 50cm$ length and $\sim 5cm$ width [11], we calculated a conservative estimate on the number of surface ossicles to be about 50 million. This estimate is based on a scale factor of 250000 for the ratio of cylindrical surface area between the adult and the juvenile. This number is very similar to that of 20 million calculated by Hampton et al. [12] for *Holothuria impatiens*.

#### List of supplementary videos

1. **Supplementary Video 1 (SV1):** Juvenile *A. parvimensis* as a system for studying cellular construction of biomineralized ossicles

2. **Supplementary Video 2 (SV2):** MicroCT fly through of discrete distributed skeleton of a juvenile *A. parvimensis*

3. **Supplementary Video 3 (SV3):** Direct imaging of ossicle morphogenesis in live juvenile *A. parvimensis*

4. **Supplementary Video 4 (SV4):** Extracellular matrix niche surrounding growing ossicles in juvenile A. parvimensis

5. **Supplementary Video 5 (SV5):** Local sclerocyte syncytial niche associated with ossicles during growth in juvenile *A. parvimensis*

6. **Supplementary Video 6 (SV6):** Surface localized cargo transport and tip localized filopodial activity in juvenile *A. parvimensis* ossicles

7. **Supplementary Video 7 (SV7):** Cytoskeletal distribution within sclerocyte syncytium in juvenile A. parvimensis ossicles

### Supplementary Methods

#### Miscellaneous methods

##### Growth series analysis for Figure 3J,K

Brightfield images were manually segmented to create binary masks representing ossicle for different stages of growth in the time series. The binary masks were skeletonized and their medial thickness calculated at each point. The 3D volume of the ossicle was approximated as a union of volumes of 3D cylinders that have lengths as distance between consecutive pixels on the skeleton and diameter as the average medial thickness of the two pixels. Surface area was calculated as the union of curved surface areas of cylinders identified similarly.

##### Vesicle tracking for Figure 5H,I

Manual tracking of two boluses represented with square and circle was performed on scale-invariant feature transform (SIFT) aligned image stacks to counter the translational motion in live video using ImageJ [13]. A custom Python code was used to calculate and plot the velocities from the raw position data.

##### Precipitation induced decoration of ossicles

The protocol to grow calcite crystallites epitaxially on top of isolated ossicles was adopted from Okazaki et al. [14]. Ossicles were isolated from small tissue sections of an adult *A. parvimensis* by digesting them in sodium hypochlorite (Clorox bleach), which dissolved all the organic tissue neighboring the ossicles, and resulted in a dense sediment containing hundreds of ossicles. The excess bleach was decanted and the sedimented ossicles were washed with distilled water with pH 8.0 (adjusted using $NaOH$). The ossicles were then dehydrated into ethanol using serial dilution (1:1, 3:1, 9:1, 1:0 ethanol to distilled water ratios) and stored. For decoration, ossicles were rehydrated into distilled water (pH 8.0) and immediately transferred into $5ml$ $0.1M$ $NaHCO_3$. To this, $2ml$ of $0.1M$ $CaCl_2$ solution was added to induce precipitaion of $CaCO_3$. After 5 minutes, the samples were then washed with distilled water (pH 8.0) and dehydrated into ethanol through a serial dilution. For visualizing crystal orientation beneath the ossicle surface, some ossicles were slightly squished under a coverslip to break several branches and expose their cross sections, before inducing precipitaation on them.

##### Scanning electron microscopy

Dehydrated decorated or undecorated ossicles we spread on one of the sticky surface of a carbon tape stuck on an SEM aluminium stub and allowed to air dry. The samples were then coated with $\sim$10 nm Au/Pd (60:40 ratio) (Denton Desk II sputter coater). Imaging was then performed in a variable-pressure scanning electron microscope (Hitachi S-3400N) operated at 5 keV.

#### Analysis of micro-CT datasets

##### Segmentation and volume and surface area calculation

The reconstructed dataset obtained as final output from the Zeiss Xradia Scout and Scan Control System software from micro-CT machine were loaded onto ORS Dragonfly Pro (version 2022.2.0.1399) software. The boundaries of the uploading region were trimmed to only import the relevant region. In case of data with multiple animals in the same volume, each animal was separated from the others by using a cylindrical volume crop. The threshold intensity range for segmenting the mineral volume was identified using the LUT slider and same values were used to generate a threshold based volume segmentation. The segmented whole animal volume was further divided into multiple ossicles volumes using a combination of seed based watershed segmentation and manual voxel removal to separate ossicles touching each other. The separated voxel groups representing individual ossicles were then converted into smooth and closed triangular meshes and exported. The volume and surface areas of these meshes were calculated using in-built functions in Pyvista library [15]. All 3D plots with colormaps and renderings of different orientations were also generated using the Pyvista library.

##### Skeletonization and cleaning

The mesh volumes were skeletonized into linear chains of voxel elements that are centrally located (medial axis) across the branches of the ossicle, using a custom built Python code inspired from a study on coral morphometrics by Ramírez-Portilla et al. [16] that utilizes ITK library [17]. The local medial thickness at each point on the skeleton was calculated as the diameter of the largest sphere that can fit within the mesh while its center lies on the skeleton point. The skeletons were stored as vtk polydata objects with identified continuous linear elements between nodes with degrees 3 or more, and medial thickness values stored as scalars on points. Linear elements with length less than 15 voxels were trimmed to get rid of false branches.

##### Conversion into graphs and simplification

Each skeleton was converted into a NetworkX [18] graph object, which we call full-graph, by assuming skeleton voxel centers as nodes and edges representing contact between voxels. The graphs were further simplified, and called simple-graph, by assuming direct edges between topologically relevant nodes of degrees not equal to 2 and medial thickness values were stored within edges as an average for all the points representing path between two nodes. An additional node, the midpoint of the thickest central edge, was identified as the origin and added to the graph.

##### Alignment, base extraction and surface fitting

For each ossicle graph, a vertical vector representing a direction perpendicular to the base of the mesh was identified using PCA over the point cloud representing the nodes from entire simple-graph. All the meshes and graphs were rotated to align the vertical direction towards positive z axis before performing further computations. The vertical portions of the pillared tables were cut off using a threshold length, resulting in graphs that represented the topology of the base of ossicles. In order to approximate the surface representing the curved base of table ossicles, we utilized base full-graphs. New nodes were interpolated between the existing nodes in the graph to fill the holes of ossicle with points. Vedo library [19] was then utilized to reconstruct and fit a surface using the interpolated points and convert it into a mesh. The resulting mesh was further remeshed, smoothened and exported for further analysis.

#### Computation of geometrical and topological quantities

##### Curvature calculations

The local principal curvatures $(p_1, p_2)$ were calculated at each point with a neighborhood radius of single average edge lengths on the surface mesh using libigl library [20]. The local mean $(\frac{p_1+p_2}{2})$ and gaussian curvatures $(p_1 \times p_2)$ were further evaluated and plotted on surfaces in 3D. The averages of mean curvature values were evaluated using a separate calculation of principle curvatures with neighborhood radius of 50 average edge lengths and then taking the numerical average over all the points of the mesh (Figure 2E).

##### Topological depth maps

For each node in the simple-graphs of the ossicles, a node's depth was evaluated as the number of edges in the shortest path connecting the origin and the node, The shortest paths connecting origin and any other node on the graph were evaluated using the djikstra algorithm. The depth of each edge were then assigned as the lower node depth amongst the depths of nodes comprising the edge. With node depths assigned, each edge was associated with a direction, pointing from the lower depth node to the higher one. Most of the nodes have one incoming and two outgoing edges associated to them, while a few of them have two incoming and one outgoing edge. The former nodes can be potential sites for bifurcation of tips, whereas the latter nodes can be used to identify the potential sites of budding (Figure S2C,D). The node depth maps were then constructed by arranging nodes with identical depth values in single rows and connecting arrows across node levels representing directional edges (Figure 2G,H).

##### Geodesic distances

The geodesic distances of all points on the ossicle mesh form a point on the mesh closest to the origin, near the base, were calculated using the pygeodesic library [21] that directly implements the exact geodesic calculation algorithm for triangular meshes [22]. The difference between the geodesic and radial planar distances were calculated and plotted on the ossicle meshes (Figure 2I).

##### Hole areas and number of sides of hole polygons

The nodes representing distinct holes were identified by calculating the minimum cycle basis for the base-graph, which creates a set of minimum number of cyclic subgraphs, that can be combined to create the entire graph. The subgraphs represent the holes and were directly used for calculating the number of sides of the polygons. To increase the accuracy for calculation of hole areas, a base graph including additional mid points for all edges was utilized to calculate the minimum cycle basis. Thus obtained cyclic subgraphs were used as marker points to cut the base surface mesh previously calculated into submeshes representing holes. The hole areas were calculated as the surface areas of these submeshes. The boundary edges of the base graphs were identified as edges which were part of only one cycle from the minimum cycle basis. The boundary nodes were identified as nodes composing the boundary edges. The identity as a boundary node or edge were stored in graphs as node and edge attribute.

##### Bifurcation angles, lengths and thicknesses

Assuming that structures grow outward from the origin, branch splitting directions can be identified for bifurcating nodes as discussed in previous sections. The angle between outgoing edges at bifurcating nodes, which are oriented in 3D space, can be evaluated computationally using vector algebra. This angle was used to represent the bifurcation angle at the nodes. Lengths were evaluated directly as the 3D separation between the nodes comprising edges. The local medial thicknesses at each voxel on the skeleton were evaluated as discussed in previous sections. For Figure 2M, the average medial thickness values were projected on to all the points on the ossicle mesh from the nearest 100 voxels from the skeleton. The thickness values were then plotted as scalar maps.

**Foot pad ossicle data**

Since foot pad ossicles in juveniles are very small due to the small size of the foot, we decided to utilize foot pad ossicles from adult *A. parvimensis* for geometrical and topological comparisons. For this purpose, foot pad tissue from adults were dissected and treated with bleach to get rid of all the soft tissue, before imaging the remaining ossicle with DIC microscopy. The observed micrograph image was then segmented, artefacts corrected and then skeletonized with medial thicknesses calculated at each point using 'medial_axis' function in skimage library [23]. The skeletons were then turned into grpahs and further geometrical and topological computations were performed.
